# 4D Single-Cell Spatial Transcriptomics Reveals Dynamic Morphogenetic Gradients and Regenerative Domains in Planarians

**DOI:** 10.64898/2026.02.18.706529

**Authors:** Kai Han, Yue Chen, Yao Li, Lidong Guo, Yuxiaofei Wang, Xiawei Liu, Yaru Lin, Zhi Huang, Qun Liu, Wenjie Guo, Rui Zhang, Wandong Zhao, Langchao Liang, Xiaoyu Wei, Li Zhou, Xuebin Mao, Jiaqi Wang, Weijian Wu, Hongwei Pan, Tao Yang, He Zhang, Xiaoshan Su, Shanshan Liu, Wenwei Zhang, Longqi Liu, Søren Tvorup Christensen, Jifeng Fei, Xin Liu, Guangyi Fan, Hanbo Li, Ying Gu, Jian Wang, Huanming Yang, Gang Pei, Xun Xu, An Zeng, Mengyang Xu

## Abstract

Regeneration relies on precise spatiotemporal gene expression and cellular responses to establish tissue identity and body patterning. Using high-resolution Stereo-seq (715 nm) on 353 sections from 16 whole animals at 8 regeneration timepoints, we constructed a 4D spatiotemporal transcriptomic map of planarian regeneration. Our analysis captured 36 refined cell types from 3,508,004 segmented cells, enabling genome-wide transcriptional imputation of gene expression dynamics across body axes at cellular, tissue, and organismal scales. We identified dynamic positional gradients and distinct spatially distributed cell types during regeneration, including an injury-induced Anterior Regenerative Zone (ARZ). The ARZ exhibited enriched positional signals in epidermal, muscle, and neural cells and was regulated by Mediator 8, which is crucial for polarity remodeling and blastema formation. This study provides a comprehensive spatial molecular and cellular map of regenerative processes, highlighting injury-induced spatial domains and key regulatory factors in planarian regeneration. We also provide an interactive web portal, offering a valuable resource for exploring and analyzing regeneration mechanisms in a spatiotemporal context.

## Introduction

Understanding the mechanisms of tissue and organ regeneration following injury is a fundamental biological question with profound implications for regenerative medicine, injury repair, and aging. Regeneration involves a complex series of events, including local and systemic responses to injury, the restoration of positional information, and the formation of new tissue structures (*1, 2*). While progress has been made in understanding these processes, several critical questions remain (*3, 4*). How do cells spatially respond to injury, and how do molecular gradients influence tissue patterning? What roles do specific cell types, morphogen gradients, and injury-responsive regions play in regeneration across complex organisms? Furthermore, how can we capture and quantify these processes at the molecular level across an entire organism?

To address these questions, profiling organisms, cells, and genes across multiple spatial and temporal scales is crucial (*5*). However, capturing continuous positional signals at molecular, cellular, and organismal levels—especially in three-dimensional space and time—remains a significant challenge (*6*). The complexity of tissue heterogeneity, large body sizes, and the preservation of cellular and molecular organization in extensive tissue sections complicate spatial transcriptomic analyses. Additionally, the absence of quantitative assays and frameworks to capture continuous positional signals at the transcriptome level in three-dimensional space and time presents an additional obstacle. This challenge is further compounded by the limited availability of classical model organisms that can fully regenerate their bodies. To date, a comprehensive three-dimensional molecular reconstruction of cellular architecture and morphogen gradients across an entire organism has not been achieved, limiting our understanding of the dynamic changes in cellular and molecular identities during regeneration.

Planarians, renowned for their exceptional regenerative abilities, serve as an ideal model for studying the spatial and temporal dynamics of tissue regeneration (*3*). These basal bilateral metazoans possess a complex anatomy (*7*), including a brain, nerve cords, peripheral nervous system, epidermis, intestine, muscles, excretory system, and a centrally located pharynx. Composed of a variety of cell types derived from three germ layers, planarians rely on pluripotent stem cells to generate and maintain their tissues (*8, 9*). They utilize precise positional cues to guide body axis formation and tissue patterning (*10, 11*). Numerous genes involved in signaling pathways for body plan patterning have been identified, expressed in a complex spatial map across the anterior-posterior, medial-lateral, and dorsal-ventral axes (*12–15*). These genes, known as position control genes (PCGs), are largely expressed in muscle tissue and play a critical role in regulating positional information during regeneration (*16*). However, the precise phenotypic outcomes associated with many of these genes remain poorly understood, and it is still unclear whether positional information is confined exclusively to muscle tissue (*16*). The regenerative process requires cells to establish, record, and interpret positional information in order to rebuild the body’s complex structure. This highlights the importance of profiling genes and cell types across multiple spatial and temporal scales. Although advances in single-cell RNA sequencing (scRNA-seq) (*17–23*) and spatial transcriptomics (ST) (*24, 25*) have enabled the profiling of cell types and gene expression patterns, these technologies still face limitations in achieving high spatial resolution at both the cellular and organismal levels. Furthermore, the mechanisms by which injury-induced local signals guide stem cells to reconstruct the body axis and regenerate fully functional three-dimensional structures remain poorly understood (*3, 26*). Thus, there is a need for comprehensive approaches that can capture the dynamic molecular and cellular events of regeneration across entire organisms.

In this study, we developed a 4D spatial transcriptomic (4D-ST) framework using the Stereo-seq technique (*27*) to create an extensive atlas of 3,508,004 segmented cells from 353 sections of 16 complete planarians, spanning eight time points of whole-body regeneration. With a resolution of 715 nm, we generated detailed transcriptional and anatomical maps of the regeneration process, constructing positional and transcriptional gradients along the body axis. Our 4D atlas, annotated with 36 refined cell types, provides a comprehensive view of gene expression dynamics across cellular, tissue, and organismal scales. Our findings offer key insights into the regenerative process. First, we reveal complete spatial gene expression patterns that define positional gradients along the body axes, tracking their dynamic changes during regeneration. Additionally, we identify an injury-induced Anterior Regenerative Zone (ARZ), marked by ROD1, which exhibits enriched positional signals in epidermal, muscle, and neural cells. The ARZ is regulated by Mediator 8 (Med8), which is crucial for polarity remodeling, blastema formation, and overall regeneration. These results provide a comprehensive molecular and spatial map of regenerative processes, highlighting dynamic changes in regeneration-responsive cells, spatial domains, and key regulatory factors.

## Results

### Building a 4D spatiotemporal transcriptomics atlas with spatial and cell-type resolution

To investigate the cellular and molecular dynamics of regeneration, we developed a framework combining spatial transcriptomics with 3D reconstruction techniques to generate a comprehensive 4D atlas of gene expression and cellular changes during the regeneration process (Han et al., submitted). We focused on pre-pharyngeal amputations, which regenerate the head, tail, and pharynx over a two-week period (Fig. 1A). Using the Stereo-seq technique (*27*), which integrates tissue cryo-sectioning with *in situ* RNA sequencing at 715 nm resolution, we profiled gene expressions at multiple stages of regeneration. Animals were collected at eight distinct time points—0 hours, 12 hours, 36 hours, 3 days, 5 days, 7 days, 10 days, and 14 days post-amputation (dpa). For each time point, two animals were sampled, resulting in a total of 16 regenerating animals. These animals were sectioned along the dorsal-ventral axis to capture the entire organism (fig. S1A). The 16 animals were processed into 10 μm thick sections, producing a total of 353 slices for spatial transcriptomics analysis using the Stereo-seq platform (Fig. 1A and fig. S1A).

**Fig. 1.**
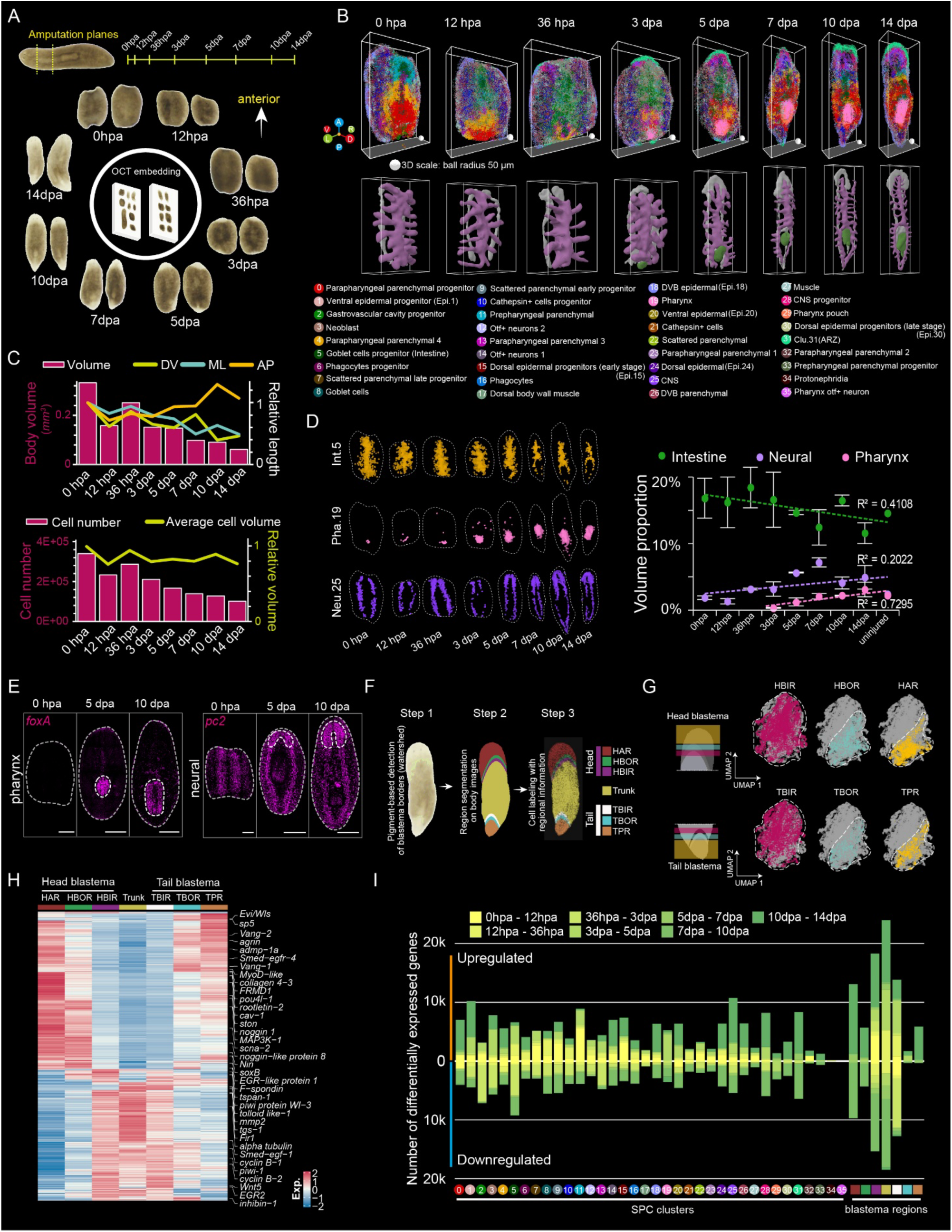
3D molecular reconstruction of whole-body planarian regeneration using 4D spatial transcriptomics. (A) Schematic representation of the amputation strategy and sampling design for planarian whole-body regeneration (WBR). The dotted line indicates the amputated pre-pharyngeal fragments. The central diagram illustrates the arrangement of 17 embedded tissue samples per block, including one intact sample and two replicates at each of the eight time points post-amputation. (B) 3D spatial visualization of 36 SPC clusters (top) and tissue meshes (bottom) in representative animals at eight time points during WBR. SPC clusters are labeled in the bottom panel. Tissue meshes highlight the intestine (purple), pharynx (green), and central nervous system (gray). (C) Top: Bar plots showing changes in body volume size; line charts depicting variations in the length of the D/V, M/L, and A/P axes relative to the 0 hpa sample at eight time points of regeneration. Bottom: Bar plots showing SPC cell counts and line charts illustrating the average cell volume across the eight time points of regeneration. (D) Left: Spatial patterns of the intestine (Int.5), pharynx (Pha.19) and CNS (Neu.25) clusters at eight regenerative time points. Right: Line plots showing the volume proportions of the reconstructed organs (Intestine, neural and pharynx) during regeneration. Error bars represent the standard deviation from two replicates. Linear fitting lines show nearly linear time-dependent changes. (E) FISH staining showing spatial patterns of pharynx (*foxA*) and neural (*pc2*) markers during regeneration. Scale bars: 500 µm. n ≥ 3. (F) Schematic diagrams illustrating the three-step process for identifying blastema subdomains based on the watershed algorithm for pigment variations. The head blastema region is divided into HAR (head anterior region), HBOR (head border outer region), and HBIR (head border inner region), while the tail blastema region includes TPR (tail posterior region), TBOR (tail border outer region), and TBIR (tail border inner region). (G) UMAP representation of cell populations from the head and tail blastema shown in (F), with distinct regions color-coded to indicate separation. (H) Heatmap depicting the relative expression of genes enriched in different blastema regions across all time points. (I) Stacked bar plot illustrating the number of differentially expressed genes in each SPC cell type or blastema region under the indicated comparison conditions.

To facilitate the generation of a comprehensive 3D reconstruction, we aligned and stitched the individual tissue sections using a newly developed 3D analysis pipeline (Materials and Methods). This resulted in 3,508,004 segmented cells across the 16 reconstructed animals, with cell counts ranging from 58,450 to 432,197 per time point after quality control (UMIs per cell >50) (Fig. 1B and fig. S1A) (data S1). This 3D-spatial transcriptomics (3D-ST) atlas spans all eight time points, providing a high-resolution, 4D view of regenerating planarians at subcellular resolution (Fig. 1B and fig. S1B). The dataset allows for the tracking of the spatial dynamics of regeneration-responsive genes and cellular interactions at various stages of regeneration. The dataset is publicly available via our searchable browser: https://db.cngb.org/stomics/prista4d (Fig. S1C).

To enhance the identification of biologically relevant tissue domains, we combined gene expression data with spatial information using a novel spatial proximity-based clustering (SPC) method. This method groups cells based on both transcriptional similarity and spatial proximity (Materials and Methods). Using SPC, we identified 36 distinct cell clusters (Fig. 1B), each characterized by unique transcriptomic profiles and spatial distributions (fig S2, A and B). For example, our analysis revealed significant spatial heterogeneity in the parenchyma, which was subdivided into 11 distinct subclusters (fig. S2A), consistent with previously identified heterogeneity (*22*). We also identified well-known tissue sub-populations, including epidermal progenitors, dorsal and ventral epidermal populations, and sub-populations of goblet cells and phagocytes within the intestine (fig. S2B). Furthermore, we discovered a new spatially localized domain, Clu.31, within the blastema. Initially emerging in both the head and tail blastemas at 36 hours post-injury, this domain eventually became restricted to the head by the end of regeneration (fig. S2A), which we designated as the Anterior Regenerative Zone (ARZ).

For better tissue contour characterization, we generated tissue meshes for the intestines, pharynx, and neuronal regions. The spatial distributions of these clusters and tissue meshes were reproducible across two animals at each time point (Fig. 1B and fig. S1B), confirming the robustness of our 3D-ST data at the organismal level. Together, this 4D-ST atlas provides a valuable resource for studying the temporal and spatial dynamics of gene expression and cellular coordination during whole-body regeneration.

### Capturing tissue and organ remodeling and identifying genes responsive to regeneration

The comprehensive 3D reconstruction enabled precise measurements of tissue volume changes through the 4D-ST dataset. Analyzing the length ratios along the dorsal/ventral (D/V), medial/lateral (M/L), and anterior/posterior (A/P) axes revealed that the D/V and M/L axes shortened, while the A/P axis elongated during regeneration (Fig. 1C, top). Although the cell count decreased, the average cell volume remained largely unchanged, suggesting that the observed volume changes were primarily due to a reduction in cell numbers (Fig. 1C, bottom). This finding aligns with the body-wide plasticity previously observed in planarians (*28*).

Using the 4D-ST dataset, we tracked the regeneration of the pharynx, as well as the remodeling of the intestine and nervous system (Fig. 1D), with validation through fluorescent in situ hybridization (FISH) for pharyngeal (*foxA*) and neural markers (*pc2*) (Fig. 1E). Both the pharynx and central nervous system, particularly the cephalic ganglia, exhibited increased volume over time, while the intestine showed a decrease in size but underwent remodeling. The pharynx began to form between 3 and 5 dpa, while the central nervous system matured by 5 dpa (Fig. 1, D and E), reflecting the gradual remodeling in these organs.

To investigate the dynamic cellular responses during regeneration, we analyzed the proportions of different cell types over time. We classified cell cluster dynamics into five patterns: continuous increase, initial increase followed by decrease, continuous decrease, initial decrease followed by increase, and unchanged (fig. S2C). Notably, dorsal epidermal progenitors (Epi.30), neural progenitors (Neu.28), and pharyngeal lineages (Pha.19 and Pha.29) exhibited a gradual increase (fig. S2, C–D), indicating an active wound response or the generation of new tissue. These results suggest that planarians may balance cell and organ proportions during regeneration, dynamically rescaling body proportions and axial polarity.

Due to the fragility of blastema tissue, conventional FISH methods are challenging for capturing internal gene expression and spatial distribution in this region (*29*). To overcome this limitation, we hypothesized that digital segmentation of the blastema region, based on pigmentation intensity, would provide a more effective means of analyzing gene expression and cellular composition. Using image recognition algorithms, we segmented the animals into head blastema, tail blastema, and pre-existing trunk regions (fig. S2E). Statistical analysis revealed positional heterogeneity in cellular responses. For instance, the pharynx (Pha.19) initially localized to the tail blastema and later to the trunk, suggesting its movement toward the body’s center. Additionally, the ARZ (Clu.31) was induced in both blastemas but persisted only in the head region (fig. S2F). This segmentation highlights the potential of our virtual 4D-ST data in identifying distinct cell subtypes that emerge at various stages and locations during regeneration.

To further investigate the molecular responses in finer regions, we divided the regenerating head and tail blastemas into three subdomains—proximal, middle, and distal—along with the trunk, and identified region-specific gene expression patterns through clustering (Fig. 1, F and G). Both the head and tail blastema exhibited similar wound-healing and remodeling gene expression, with enrichment in genes associated with Wnt and BMP signaling pathways. However, different subdomains within these regions showed distinct gene expression profiles (Fig. 1H, data S2). Additionally, comparison of differentially expressed genes within the same clusters or regions across various regeneration time points revealed temporal variation in gene expression patterns. For example, we observed that parenchymal cell types and epidermal progenitors responded to injury within the first 12 hours post-amputation (hpa), while goblet cells and *cathepsin*^+^ cells were activated between 12 hpa and 3 dpa. Neuronal cells and the ARZ (Clu.31) domain showed a response after 3 dpa (Fig. 1I, data S2), highlighting distinct cellular and regional responses at different stages of regeneration.

In summary, the 4D-ST atlas of whole-body regeneration offers an in-depth view of tissue remodeling, dynamic changes in cellular localization, and the spatial distribution of cell populations throughout planarian regeneration.

### Spatiotemporal dynamics of positional gradients during whole-body regeneration

In an accompanying study, we identified genes with regional expression patterns in 3D intact planarians, which we proposed as spatially biased genes (SBGs) (Han et al., submitted). Some of these genes, involved in patterning processes, were classified as positional control genes (PCGs) (*10*). To investigate the dynamics of these SBGs during regeneration, we employed 4D-ST to map gene expression across the entire organism. We hypothesized that injury would disrupt SBG expression, with recovery occurring gradually as regeneration progressed.

To test this, we analyzed the spatiotemporal patterns of SBGs by mapping representative samples from each time point onto clusters along the A/P axis using logistic regression (Fig. 2A). Our results showed that, following injury, most SBG clusters were initially disrupted and then fluctuated like waves along the body axis. Over time, these patterns gradually stabilized, resembling the uninjured state, though at different rates. By 5 dpa, the overall gradient patterns had largely been restored to resemble those of uninjured individuals (Fig. 2A), highlighting the dynamic process of positional remodeling during regeneration.

**Fig. 2.**
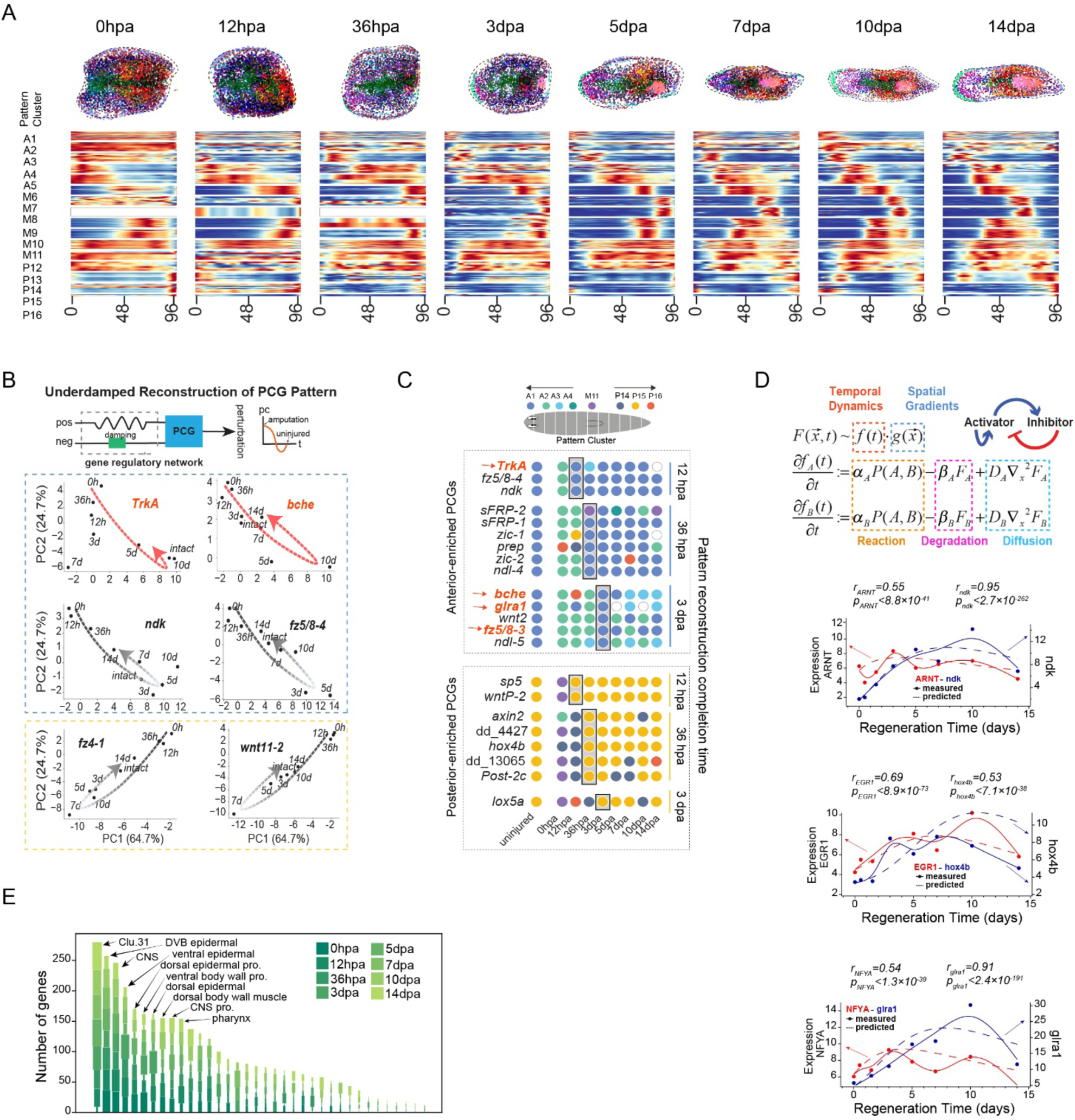
Dynamics of spatially biased genes (SBGs) during whole-body regeneration. **(A)** Heatmaps showing changes in spatial expression patterns along the A/P axis during regeneration for the 16 A/P pattern cluster genes. These genes exhibit spatially biased expression along the A/P axis in intact animals. Hollow rectangles indicate the absence of specific gene expression clusters at particular time points. Animals are virtually divided into 100 sections along the A/P axis. Left margin annotations indicate cluster numbers. **(B)** PCA representation of pattern reconstruction for known and potential PCG candidates. PC1 corresponds to the head-tail gradient feature, while PC2 captures fluctuations in the pharynx region (convex and concave). Time-resolved trajectories in PCA space reveal universal self-organized dynamics during SBG pattern regeneration, resembling an underdamped mass-spring system (top). **(C)** Schematic diagrams illustrating the hierarchical reconstruction of SBG patterns, colored by cluster ID. SBGs are arranged by the timing of repatterning completion, as shown on the right. Hollow circles indicate the absence of expression at specific time points. Red arrows highlight newly identified SBGs. The color gradient represents the recovery of SBGs at different stages post-amputation. The box highlights the time points when repatterning is complete. **(D)** Spatiotemporal modeling of SBG patterns during regeneration. The interaction between SBGs and their upstream regulators generates Turing patterns in an autoregulatory activator-inhibitor system (top). Gene expression at any time point after amputation can be predicted. The lower line charts compare predicted repatterning (dashed line) with measured data (solid line) for both known and newly identified SBGs. r, Pearson’s correlation. **(E)** Bar plot showing the number of SBGs highly enriched in each SPC cluster at different time points during regeneration.

Leveraging the quantitative nature of spatial transcriptomics, we further explored the spatiotemporal dynamics of SBGs. Principal component analysis (PCA) on known PCGs along the body axes of intact planarians revealed that genes expressed in the same regions clustered together. For example, *ndl-4* and *sfrp1* were specifically expressed in the head (fig. S3A), confirming their region-specific patterns (*30, 31*). This analysis also allowed us to quantify relative expression patterns, such as *ndk* in both the head and pharynx, and *wntA* in the pharynx (Fig. S3B), consistent with established spatial distributions (*14, 32–34*).

Next, we mapped SBGs across multiple time points in PCA space to track their dynamic recovery during regeneration. Notably, several known PCGs, including *ndk, fz5/8-4*, *fz4-1*, and *wnt11-2* (*14, 32*), exhibited a reciprocal recovery pattern along the A/P axis. These genes initially showed higher expression compared to uninjured individuals, before gradually returning to baseline levels (Fig. 2B, bottom). Following amputation, the disruption of PCG expression resembled a perturbation in a dynamic system, akin to an underdamped mass-spring system (*35*), where the system is displaced from equilibrium, oscillates, and then stabilizes. In line with this, some SBGs displayed wave-like expression along the A/P axis. This dynamic recovery trajectory eventually restored the gene expression profile to its homeostatic state at 14dpa (Fig. 2A). We hypothesize that injury disrupts regional expression, and that the gene regulatory network may act as a restoring force, guiding the recovery of disrupted PCG expression through feedback mechanisms (Fig. 2B, top). Consistent with this model, our analysis showed that PCG expression was elevated between 3 and 10 dpa, and returned to baseline by 14 dpa upon completion of regeneration (Fig. S3C), further supporting this recovery model.

To visualize the dynamic recovery of spatial patterns, we color-coded clusters of known PCGs at each time point (Materials and Methods). These patterns were restored by 12 hours, 36 hours, or 3 dpa (Fig. 2C), suggesting a temporal transformation in their recovery. Notably, spatial patterns were restored earlier than detectable changes in gene expression. For instance, the spatial pattern of *ndk* was restored by 12 hpa (Fig. 2C), while its expression began to increase only at 36 hpa (Fig. S3C, data S3). This temporal dynamic suggests that spatial patterning may influence the regulation of gene expression.

Motivated by the underdamped response of PCG trajectories (Fig. 2B) and the temporal hierarchy between spatial pattern recovery and gene expression (Fig. 2C), we hypothesized that SBG expression dynamics could be modeled mathematically. Extending the self-organizing model proposed for the Wnt pathway along the A/P axis (*30*), we applied the Gierer-Meinhardt model of a simple activator-inhibitor system (*36*) within the Turing system framework (*37*) to predict global changes in SBG expression during regeneration. By analyzing spatial gradients in exponential form and excluding the influence of the pharynx-enriched genes, we separated temporal and spatial components to simulate changes in activator and inhibitor expression at each time point during regeneration (Fig. 2D, top). The predictions from this model were largely consistent with the expression values measured in the Stereo-seq data for genes such as *ARNT* (*38*), *Ndk* (*14*), *EGR1*, *hox4b*, *Nfya*, and *glra1* (Fig. 2D, bottom), suggesting that the model offers a reasonable representation of the observed dynamics.

We next explored whether SBGs were enriched in specific cell types or regions. By quantifying the number of SBGs enriched in each SPC at different stages of regeneration, we observed that SBGs were expressed across multiple lineages, including muscle, epidermal, and neural lineages (Fig. 2E). Interestingly, the ARZ domain (Clu.31) displayed characteristics from several lineages and contained the highest number of SBGs (Fig. 2E). This observation suggests that the ARZ may play a role in remodeling and maintaining polarity. Overall, our 4D-ST analysis provides a detailed view of the spatiotemporal dynamics of SBG expression during regeneration, offering support for a model based on self-organized reaction-diffusion patterns.

### Characteristics of the injury-induced Anterior Regenerative Zone (ARZ) enriched in SBGs

Having demonstrated that the ARZ (Clu.31) exhibits injury-induced anterior localization and is enriched in SBGs, we hypothesize that the ARZ plays a crucial role in regulating PCGs and maintaining regenerative polarity during regeneration, similar to the proposed function of the organizer (*39*). We further sought to characterize this region. At homeostasis, the ARZ is localized to the anterior side, with signatures of three distinct lineages (Fig. 3A and data S5). Gene expression analysis within this domain revealed enriched expression of SMED30003831 (*smed03831 or Rod1*)*, caveolin3*, and SMED30001640 (*smed01640)* (fig. S4A and data S2). Notably, the ARZ spans both the peripheral epidermal and subepidermal areas of the head, distinguishing it from the *Equinox*-expressing wound epidermis (*40*) (fig. S4A). Gene ontology (GO) analysis of ARZ-enriched genes identified processes related to epidermal differentiation, muscle contraction, and neural development (fig. S4B). Co-FISH experiments with ARZ marker *smed03831* and lineage markers confirmed that the ARZ encompasses epidermal (*agat-1*) (Fig. 3B), muscular (*collagen*) (Fig. 3C), and neural (*pds*) cells (Fig. 3D), solidifying its tri-lineage characteristics. These findings suggest that the ARZ is a co-regulated, regeneration-responsive region.

**Fig. 3.**
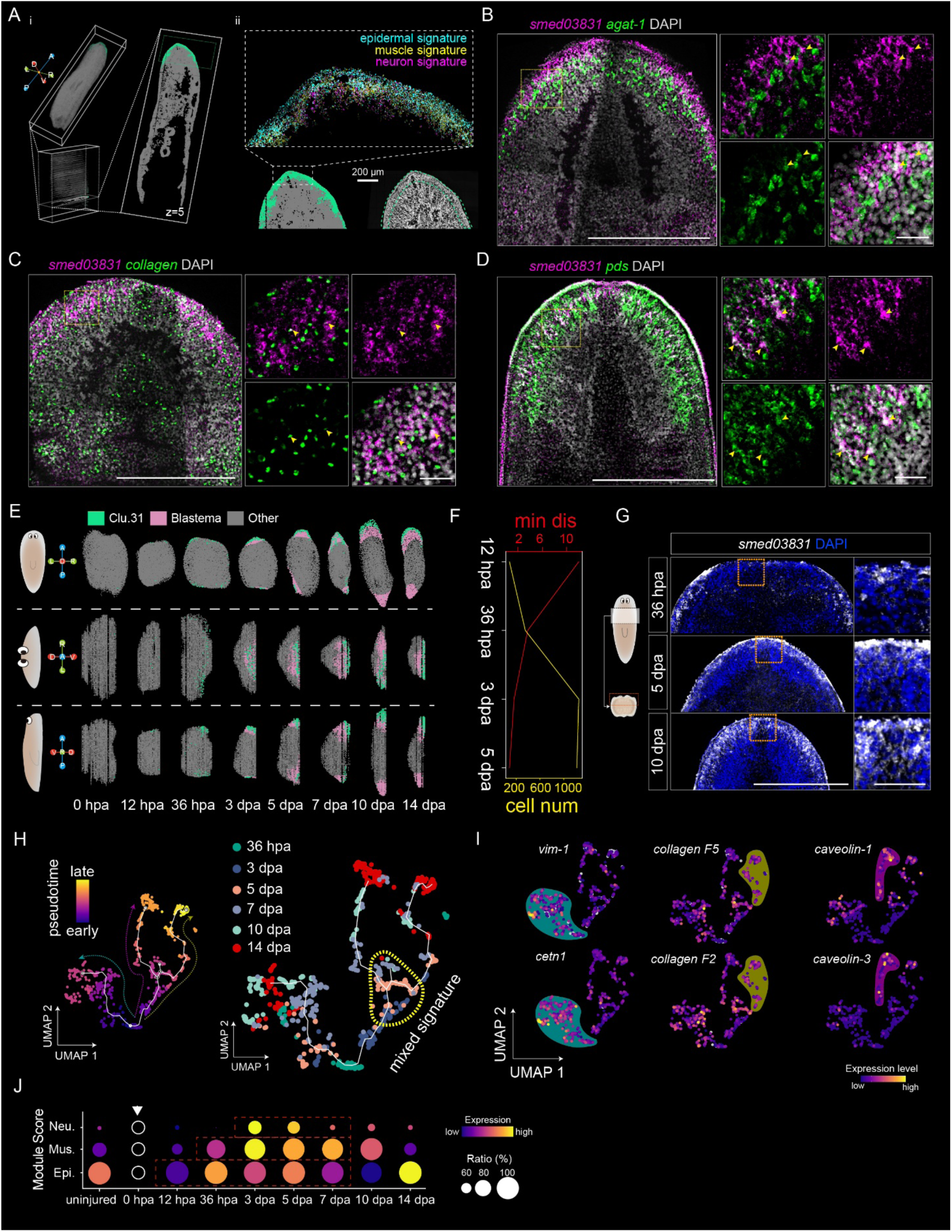
Identification and characterization of the ARZ (Clu.31) domain in the blastema region. **(A)** Spatial visualization of the Clu.31 domain in a homeostatic worm. (i) 3D spatial transcriptomics data showing Clu.31 (green) with the right panel displaying the fifth section from ventral to dorsal (z = 5). (ii) Top: Enlarged view of three lineage signatures within Clu.31 in a single slice. Bottom left: Enlarged view of the head region from (i). Bottom right: ssDNA staining highlights the presence of multiple cell layers coexisting in Clu.31, with the green dashed line indicating the location of Clu.31. **(B-D)** FISH staining for the Clu.31 marker (*smed03831*, magenta) with the epidermal marker agat-1 (green) (B), muscle marker collagen (green) (C), and neuron marker pds (green) (D). Co-expressed cells are indicated by yellow arrowheads. Scale bars: 500 µm (left); 50 µm (right). n ≥ 3. **(E)** Spatial distribution of the Clu.31 domain during regeneration, shown from top, front, and side views. Green dots represent SPC cells within Clu.31, pink dots represent SPC cells in the blastema region, and grey dots represent other SPCs. **(F)** The position and cell number of Clu.31 during regeneration. Top: Line plot (red) showing the decreased minimal distance (min dis) of Clu.31 to the wound surface, accompanied by an increase in the cell number (Cell num) of Clu.31 during wound healing (yellow line). Min dis represents the minimal distance to the wound surface in UV spatial coordinates, while Cell num refers to the number of Clu.31 cells. **(G)** FISH staining showing *smed03831* expression in the head blastema during regeneration. Enlarged areas are shown to the right. n ≥ 3. Scale bars: 500 µm (left); 50 µm (right). **(H)** Pseudotime trajectory analysis of Clu.31 across six time points of whole-body regeneration (WBR), from 36 hpa to 14 dpa. SPC cells exhibiting mixed neural, muscular, and epidermal markers are indicated by a yellow dashed circle. **(I)** Feature plot showing the expression of representative cell-type marker genes—epidermis (left), muscle (middle), and neuronal (right) lineages—along pseudotime trajectories from (H). **(J)** Bubble plot displaying the gene set module scores of markers for neural (Neu.), muscular (Mus.), and epidermal (Epi.) lineages within Clu.31 during regeneration.

To track the temporal dynamics of the ARZ, we analyzed its spatial location throughout regeneration. At 36 hpa, the ARZ was present as scattered clusters near the ventral wound sites in both head and tail fragments. By 3–5 dpa, these cells converged towards the midline and expanded to cover the wound area, coinciding with wound closure and blastema formation. By 10 dpa, ARZ cells diminished in the tail but persisted in the head region (Fig. 3E, fig. S4C). Measurement of the distance from the wound surface revealed that the ARZ gradually approached the amputation site during the first five days, with an increase in cell number within the zone (Fig. 3F). This was further confirmed by staining for the ARZ marker *smed03831* in the regenerating head region (Fig. 3G).

To investigate the putative origin of ARZ cells, we traced their pseudotime trajectory during regeneration using Monocle (*41*). This analysis revealed three distinct branches (Fig. 3H), each enriched for genes specific to epidermal, muscle, or neuronal lineages (Fig. 3I, Fig. S4D). The earliest reappearance of epidermal signatures at 12 hpa marked the emergence of the ARZ, followed by muscle and neuronal markers at 3 dpa, coinciding with blastema formation (Fig. 3J). By 14 dpa, the ARZ cellular composition had largely reverted to epidermal cells, resembling the homeostatic state (Fig. 3J). The expression of ARZ-enriched genes aligns with these cellular dynamics (data S5), further supporting the coordinated and timely assembly of the ARZ domain during regeneration.

### Cellular composition and regulation of the polarity-enriched ARZ domain

Having demonstrated the coordination between the ARZ formation and regeneration, we next sought to investigate the cellular components that control the formation of the ARZ. Our focus was on the epidermis, as it is the earliest cell type to emerge within the ARZ. To identify the regulatory factors involved in ARZ formation, we employed RNA velocity, a method that distinguishes between unspliced and spliced mRNAs (*42*), to predict the putative trajectory of the SPC clusters in epidermal lineages (fig. S5A). The velocity vectors indicated that epidermal cells in the ARZ primarily originate from the ventral epidermal lineage (Epi.1) (fig. S5, A and B). To visualize ARZ formation in a spatiotemporal context, we projected 3D spatial data onto 2D maps of the head blastema at various time points using UV mapping (Fig. 4A), a technique used in 3D modeling to apply a 2D image (texture) onto the surface of a 3D model (see Materials and Methods). Applying pseudotime trajectory analysis on the 2D map after UV mapping revealed a cell-state transition from ventral epidermal cells toward the ARZ, while left and right dorsal-ventral boundary cells on both sides of the head blastema moved toward the anterior pole (Fig. 4, B to D). These observations suggest that ARZ formation involves interactions between ventral and dorsal epidermal cells. Indeed, comparing spatial maps at 36 hpa and 3 dpa reveals the expansion of the ARZ domain from both ventral and dorsal sides, coinciding with wound closure (Fig. 4E).

**Fig. 4.**
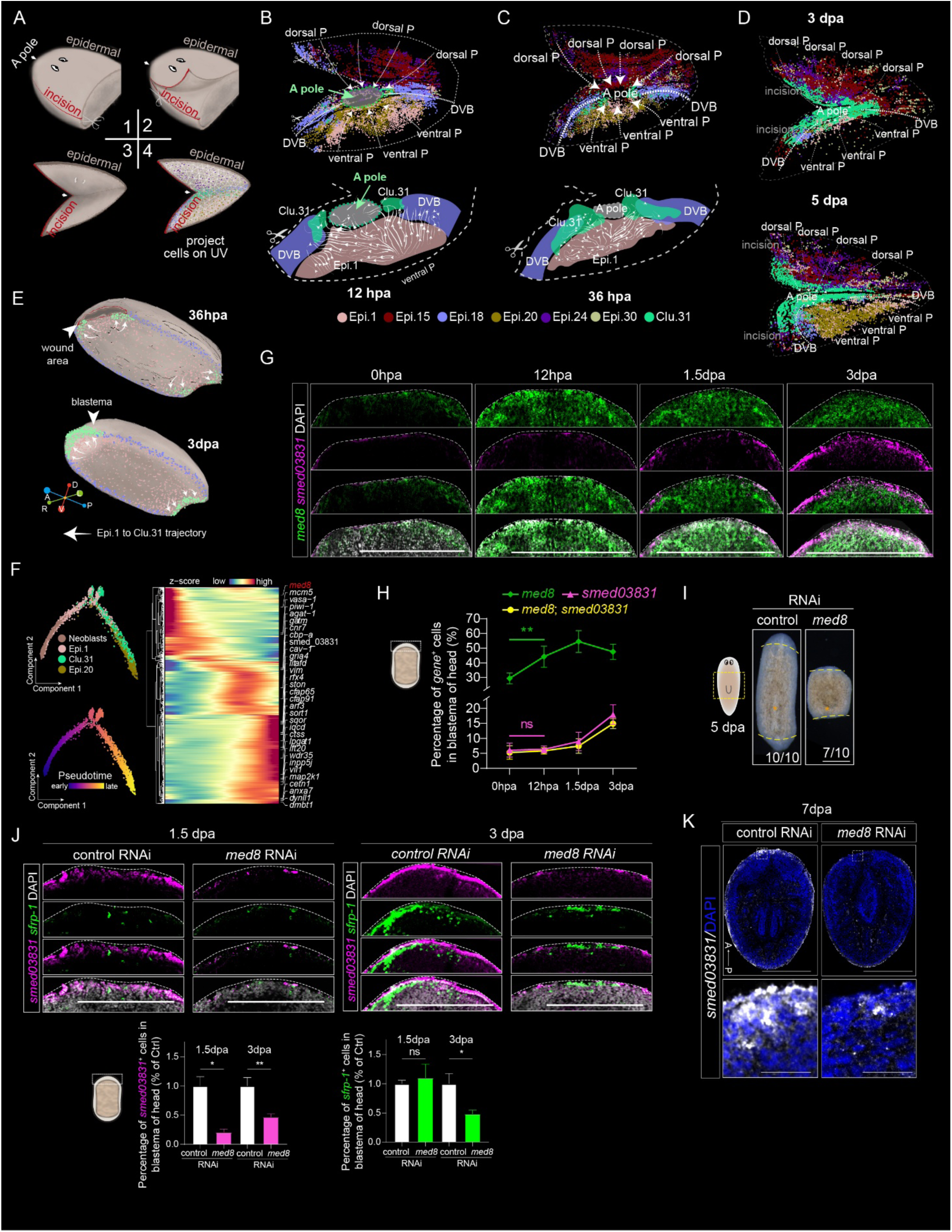
Cellular composition and regulation of the ARZ domain. **(A)** Schematic illustrating the workflow for unwrapping the 3D epidermal surface of the head blastema into 2D spatial coordinates. UV unwrapping of the 3D mesh was performed by manually marking a seam along the D/V boundary (Step 1) and unwrapping the surface by cutting along the seam (Steps 2, 3). Epidermal cells were projected onto the 2D coordinates by minimizing the distance between each 3D cell center and the nearest subdivision vertices (Step 4). See Materials and Methods for details. **(B)** UV spatial mapping of the ventral epidermal transition forming the Clu.31 domain at 12 hours post-amputation (hpa). Top: Distribution of Clu.31 (green) and epidermal SPCs on the unwrapped UV map, with dot colors representing SPC types. Labels and dashed lines indicate key positions. Bottom: Predicted cell transition streams of the epidermis as modeled using Dynamo. DVB, Dorsal-Ventral Boundary; A pole, anterior pole; P, posterior. **(C)** UV spatial map showing Clu.31 transition patterns at 36 hpa. Top: Distribution of Clu.31 and epidermal SPCs on the unwrapped UV spatial map. Bottom: Cell transition streams of the epidermis predicted using Dynamo. **(D)** UV spatial map showing Clu.31 transition patterns at 3 days post-amputation (dpa, top) and 5 dpa (bottom). The distribution of Clu.31 (green) and epidermal SPCs on the unwrapped UV spatial map is shown. **(E)** Visualization of Clu.31 movement during wound healing (top, 36 hpa) and blastema formation (bottom, 3 dpa). The white arrow indicates the predicted trajectory of the epidermal transition, delineated based on the spatial dynamo shown in (C) and (D). Arrowheads highlight the location of either the wound surface (top) or blastema (bottom). **(F)** Pseudotime trajectory analysis of ventral epidermis and neoblast cells. Left: Distinct states of SPCs identified by pseudotime analysis, with cells colored by SPC clusters (top left) and pseudotime (bottom left). Right: Heatmap showing significantly altered genes discovered by Monocle 2 along the trajectory. **(G)** Expression and localization of *med8* and *smed03831* in the head blastema of regenerative fragments at the indicated time points. Scale bars, 500 µm. **(H)** Percentage of *med8*^+^, *smed03831*^+^, or co-expressing cells in the blastema shown in (G). ns, *p* > 0.05; \*\**p* < 0.01; two-tailed unpaired *t*-test. **(I)** Representative phenotypes following *med8* RNAi at 5 dpa. n = 10 animals for each condition. Scale bars: 500 µm. **(J)** Expression and localization of *smed03831* and *sfrp-1* in the blastema of control and *med8* RNAi animals at 1.5 (top left) and 3 dpa (top right). Scale bars: 500 µm. The ratio of *smed03831*- or *sfrp-1*-expressing cells in the blastema was quantified (bottom). ns, *p* > 0.05; \**p* < 0.05; \*\**p* < 0.01; two-tailed unpaired *t*-test. **(K)** FISH staining of *smed03831* in control and *med8* RNAi animals. n ≥ 3 animals per condition. Scale bars: 500 µm (top); 50 µm (bottom).

To explore the role of the ARZ in positional control during regeneration, we conducted trajectory analysis to identify potential regulators involved in cell differentiation within the ARZ (Fig. 4F). Notably, the mediator complex subunit 8 (*med8*) emerged as a top differentially expressed gene along the pseudotime trajectory (Fig. 4F). Med8 is an essential component of the mediator complex, playing a critical role in transcription regulation (*43*). The planarian *med8* homolog is evolutionarily conserved and shares close homology with orthologs in other species (fig. S5C). In the homeostatic state, *med8* is highly expressed in neoblasts (fig. S5D), with enrichment observed across multiple neoblast subpopulations (fig. S5E-F). Following injury, *med8* expression increased in the wound area at 12 hpa, prior to the emergence of *smed03831*^+^ cells at 1.5 dpa (Fig. 4G). This suggests that *med8* may regulate ARZ formation. Co-expression of *med8* with *smed03831* in a significant portion of blastema cells after 1.5 dpa further supports its role in ARZ formation (Fig. 4, G and H). To assess the functional role of *med8*, we performed RNA interference (RNAi) knockdown experiments (fig. S5G) and measured the expression of the ARZ marker *smed03831*. Knockdown of *med8* resulted in impaired blastema regeneration by 5dpa (Fig. 4I) and a significant reduction in the number of *smed03831*^+^ cells at 3 and 5 dpa (Fig. 4J), suggesting that *med8* is required for ARZ formation.

Having shown that *med8*(RNAi) hinders ARZ reconstruction (Fig. 4J), we next investigated the specific stage at which *med8* influences ARZ formation by examining gene expression at different regeneration time points. Given that the ARZ is enriched with various PCGs (Fig. 2E), we hypothesized that *med8* knockdown might disrupt polarity remodeling. To test this, we examined the expression of the anterior pole marker *sfrp-1* (*13, 44*), which was significantly reduced upon *med8* knockdown (Fig. 4J), suggesting defects in pattern formation during regeneration. In control animals, *smed03831*^+^ cells were enriched at the wound site at 1.5 dpa and fully covered the wound, followed by the appearance of *sfrp-1*^+^ cells at the anterior pole by 3 dpa (Fig. 4J). This temporal sequence suggests that ARZ formation precedes anterior pole formation. In contrast, *med8*(RNAi) animals exhibited impaired ARZ formation and a reduction in *sfrp-1*^+^ cells (Fig. 4J), indicating a disruption in patterning. By 7 dpa, ARZ formation was completely disrupted in *med8*(RNAi) animals, and regeneration failed (Fig. 4K), linking ARZ formation to successful regeneration. Together, these findings support the idea that *med8*-mediated ARZ formation is essential for proper pole formation during regeneration.

### *Mediator 8* regulates ARZ formation to control differentiation and blastema development

To identify the transcriptional programs mediating changes in the ARZ, we conducted scRNA-seq on *med8* and control RNAi animals with amputated tails undergoing head regeneration. By integrating the data from both groups, we identified known cell lineages, including stem cell populations and eight distinct cell types (Fig. 5A), consistent with previous findings (*8, 22*). Notably, *med8* RNAi led to an increase in the neoblast population, while the proportions of lineages associated with the ARZ—specifically neural, muscle, and epidermal—were reduced (Fig. 5B). This observation was further confirmed by pseudotime trajectory analysis using Monocle, which revealed similar reductions in the differentiation of these cell lineages (fig. S6A). Label transfer analysis, performed to match ARZ cells in the scRNA-seq dataset, confirmed a decrease in cells representing each ARZ component in *med8* RNAi animals (Fig. 5C and fig. S6B).

**Fig. 5.**
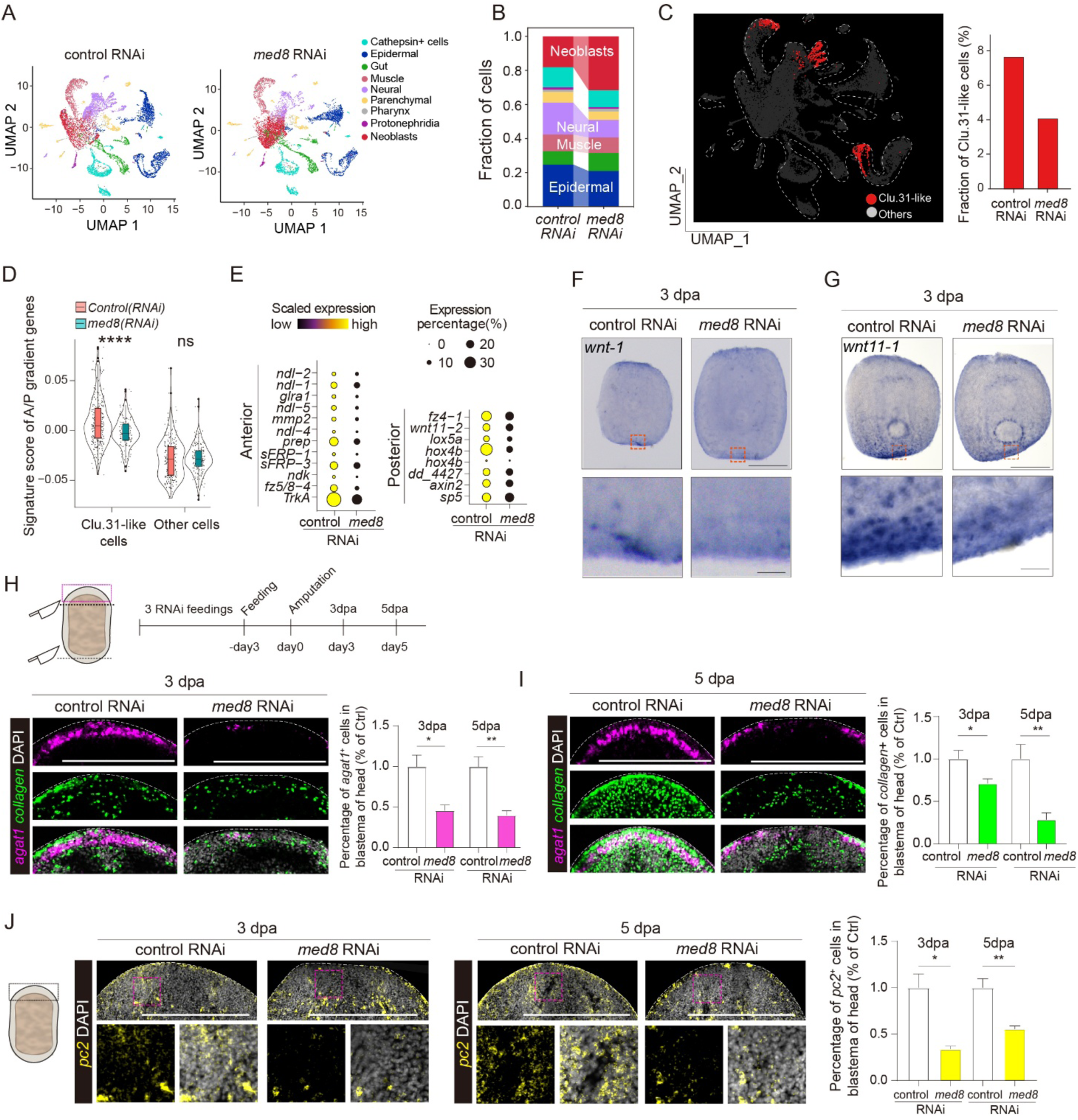
Med8-dependent regulation of anterior regenerative zone and blastema formation. **(A)** UMAP visualization of scRNA-seq analysis depicting cell lineages from control and *med8* RNAi-treated tail fragments. **(B)** Bar plot showing the changes in cell populations following med8 RNAi knockdown. **(C)** Left: UMAP visualization of neural, muscular, and epidermal cells highlighting the ARZ signature (Clu.31-like) in scRNA-seq data. Right: Bar plot illustrating the decrease in the fraction of ARZ (Clu.31) cells in *med8* RNAi-treated animals compared to controls. **(D)** Violin plot showing changes in the module score of A/P gradient genes in ARZ (Clu.31) cells. p-values are from Wilcoxon test: ns, *p* > 0.05; ****, *p* < 0.0001. **(E)** Dot plot illustrating the relative expression of representative PCGs in anterior or posterior regions from scRNA-seq data. **(F-G)** WISH analysis showing the expression and localization of posterior markers *wnt-1* (F) and *wnt11-1* (G) in control and *med8* knockdown animals. Scale bars: 500 µm (top), 50 µm (bottom). n = 6 animals with consistent results. **(H-I)** Expression and localization of muscle marker *collagen* and epidermal marker *agat-1* in the blastema of control and *med8* RNAi animals at 3 (H) and 5 days post-amputation (dpa) (I), respectively. Scale bars: 500 µm. n = 6 animals with consistent results. The percentage of positive cells in the blastema was quantified. ns, *p* > 0.05; \**p* < 0.05; \*\**p* < 0.01, two-tailed unpaired t-test. **(J)** Expression and localization of the neural marker *PC2* in the blastema of control and *med8* RNAi animals at 3 and 5dpa. Scale bars: 500 µm. The percentage of positive cells in the blastema was quantified (right). \**p* < 0.05; \*\**p* < 0.01, two-tailed unpaired t-test. All FISH images are maximum-intensity projections. Error bars represent SEM. n ≥ 3 biologically independent experiments.

Given that the ARZ is enriched for genes involved in polarity formation, we next examined whether *med8* RNAi caused global defects in the expression of anterior and posterior pole markers. Analysis of A/P axis marker gene expression in the scRNA-seq data revealed that *med8* RNAi led to a reduction in the polarity signature score within ARZ cells, but not in other cell types (Fig. 5D and fig. S6C). Further analysis showed a decrease in both anterior and posterior markers (Fig. 5E). Whole-mount in situ hybridization (WISH) confirmed reduced expression of posterior markers, including *wnt1* and *Wnt11-1* (*34*), though body-wide polarity was not completely disrupted in *med8* RNAi animals (Fig. 5, F and G), suggesting that the observed changes are linked to regenerative growth. These results indicate that *med8* RNAi impairs ARZ formation and alters global positional information within the blastema during regeneration.

To investigate how *med8* affects ARZ formation, we examined the expression of ARZ markers within the blastema. Major cell types in the ARZ, including epidermal (*agat1)* and muscle (*collagen*) cells, were reduced in *med8* RNAi animals (Fig. 5H). Additionally, the expression of transcription factors essential for the differentiation of neural (*tcf/lef-1*) (*45*), epidermal (*p53*) (*46*), and muscle (*dmrt2*) cells was diminished upon *med8* knockdown (Fig. S6D), indicating that *med8* is crucial for maintaining the transcriptional programs associated with ARZ cell fate (*47*). In line with this, gene expression analysis in Nb2 neoblasts revealed downregulation of pathways related to stem cell division and neural and epidermal fate determination (Fig. S6E). Furthermore, FISH staining at 3 dpa and 5 dpa demonstrated reduced expression of markers for epidermal and muscle cells in the blastema region (Fig. 5, H and I), suggesting impaired cell fate decisions in ARZ cells. We also observed decreased expression of neural markers in *med8* RNAi animals (Fig. 5J), further confirming the disruption of ARZ lineage specification and impaired head regeneration.

Finally, we assessed whether *Med8*-mediated ARZ formation is required for homeostasis. While short-term *med8* RNAi treatment caused minimal phenotypic changes in homeostatic animals, prolonged *med8* RNAi led to head regression (Fig. S6F), suggesting that sustained loss of ARZ function results in homeostatic defects. In summary, our data support the role of *med8*-regulated ARZ in controlling blastema growth by regulating positional information, which is essential for proper tissue regeneration and the maintenance of homeostasis.

## Discussion

Understanding the full spectrum of spatial information and the principles governing pattern formation during regeneration in tissues and organs remains a significant challenge. This complexity is driven by the intricate tissue geometry, the large size of multicellular organisms, and the limited number of model organisms capable of regenerating entire tissues. Additionally, the absence of techniques capable of capturing high-resolution spatial transcriptomic data across an entire organism in 3D over time further complicates this challenge. In this study, we applied high-resolution Stereo-seq (715 nm) to planarians to develop a framework for reconstructing the 4D spatiotemporal landscape of whole-body regeneration. Our 4D-ST dataset addresses some of the limitations of traditional techniques, such as resolution, low-throughput FISH assays, and single-slice-level spatial transcriptomics, providing a holistic and high-resolution view of genes, cell types, and spatial domains before and during regeneration. While recent spatial transcriptomics methods provide spatial context (*24, 25, 48*), they often lack full 3D or single-cell resolution, limiting the ability to comprehensively profile morphogen gradients and cell types across entire organisms over time. In contrast, our 4D regeneration atlas offers a high-resolution, time-resolved framework for analyzing regenerative dynamics. Exploring our dataset allowed for the recovery of single-cell transcriptomes at the temporospatial level, enabling the visualization of gene expression patterns, positional signals, and cell type distributions across multiple scales. This analysis revealed morphogen gradient gene dynamics, identified regenerative domains, and highlighted key regulatory factors. The full dataset is available through our online resource: https://db.cngb.org/stomics/prista4d.

By leveraging the complete repertoire of SBGs across four dimensions, we examined gene expression across body regions and scales, particularly within the delicate blastema region, allowing for quantitative analyses across multiple body regions. We confirmed known PCGs and identified potential new PCG candidates, facilitating the modeling of morphogenetic gradients using the Turing reaction-diffusion model. While the Turing system has been proposed for fission and regeneration (*30, 37, 49*), its applicability to modeling planarian regeneration remains unclear (*50*). Our real dataset supports simulation predictions, with some genes showing consistency with actual data. This rich dataset and quantitative approach provide a foundation for studying scalable, self-organized pattern formation in more detail, offering a framework for understanding how positional information is re-established during regeneration. Integrating high-resolution spatial transcriptomics with single-cell analysis, our study offers a valuable resource for investigating how positional information is maintained, disrupted, and interpreted across different tissues and cell types following injury. This dataset bridges the gap between molecular, cellular, and morphological aspects of regeneration, offering a comprehensive multimodal view of whole-body regeneration dynamics in organisms.

Muscle cells have long been recognized as the primary conveyors of positional cues in adult planarians (*16*). However, our 4D-ST dataset expands this view, identifying non-muscular lineages, such as neuronal and epidermal cells, as contributors to the regenerative positional landscape. This suggests that pattern remodeling involves a multi-lineage process, with multiple cell types participating in encoding, reading, and interpreting positional information (*51, 52*). We comprehensively profiled SBGs, which extend beyond PCGs, to define a systemic patterning system involving genome-wide spatial regulation. Since regenerative patterning involves both global polarity and local fate specification, most SBGs may not show defects in polarity but could alter cell fate (*31, 53, 54*). The temporal dynamics of different SBG classes suggest that encoding and interpreting positional gradients is a distributed and hierarchical process, coordinated among various cell types and exhibiting self-organizing properties within the organism.

Analysis of SBG distribution revealed a multi-lineage mixing domain, which we define as the Anterior Regenerative Zone (ARZ, Clu.31). This region is characterized by high expression of SBGs and dynamic changes in cellular proportions. Notably, we identified ROD1 (SMED30003831), a neuronal-enriched gene, as a marker of the ARZ, suggesting the presence of distinct cell types within this domain. The extent of the ARZ appears to surpass that of the anterior pole (*55*) , with significant changes in intrinsic cell types, indicating that it may function as a co-regulatory domain. The relationship between the ARZ and the anterior pole remains unclear, and whether the ARZ possesses inductive activity warrants further investigation (*56*). From an evolutionary perspective, the presence of this spatiotemporally conserved regenerative domain (ARZ) is reminiscent of the apical epithelial cap (AEC) observed in vertebrate appendage regeneration (*57*). Our findings suggest potential structural and functional similarities between the planarian ARZ domain and the AEC of regenerating axolotls, including the conservation of molecular marker gene expression (e.g., Wnt/β-catenin and FGF pathways) (*13, 14, 44*). In axolotls, the blastema forms beneath the thickened multilayered AEC, where cells differentiate and grow to replace lost structures (*58, 59*). While we observed parallels in blastema formation and differentiation, planarians exhibit intricate temporospatial cellular dynamics during regeneration. Further comparative and functional analyses across metazoans could help clarify whether specific positional circuits and molecular programs are evolutionarily conserved across species (*39*).

Our findings using *Med8(RNAi)* as a proxy suggest that epidermal, muscular, and neural cells within the ARZ likely contribute to positional information for blastema induction. The persistence of this domain in adult planarians may help explain their homeostatic maintenance, providing insights into the regulation of regeneration (*39, 60*). We further explore that *Med8*, a subunit of the Mediator complex acting as a bridge between transcription factors and RNA polymerase II (*43*), modulates key genes related to epidermal, muscle, and neural specification within the regenerative domain. The Mediator complex is known for its role in maintaining stem cells, as well as in lineage-specific differentiation (*47*). While the loss of Smed-*med14* specifically affects stem cell populations, the loss of *Med8* does not (*61*), suggesting that distinct Mediator components have different requirements for stem cell function in planarians. Our findings build upon previous studies and demonstrate that *Med8* is important for the differentiation of stem cells into neural, muscle, and epidermal lineages within the ARZ region, as well as for generating proper regenerative patterning signals. This suggests that Mediator, in conjunction with transcription factors (*62–67*), may be involved in establishing the epigenetic landscape necessary for regenerative patterning.

Despite the advantages of our 4D-ST approach, some limitations remain. First, the sequencing depth is lower compared to scRNA-seq, which may impact the detection of rare cell populations. Second, there are limited biological replicates due to the technical challenges of generating whole-organism 4D-ST data. Additional replicates and validation will be required to further assess the robustness of morphogenetic gradients across individuals. Future efforts should focus on increasing sequencing depth, expanding the number of biological replicates, and incorporating complementary approaches to validate the dynamics of gene regulation. Nevertheless, integrating our data with gene perturbation and longitudinal imaging studies will enable us to directly assess the functional contributions of specific positional signals in guiding pattern remodeling and regeneration.

Our study establishes a framework for a four-dimensional, high-resolution atlas of gene expression dynamics throughout whole-body regeneration. By combining spatial and temporal transcriptomic data, we provide a novel framework for understanding the principles governing regenerative patterning, advancing both regenerative biology and spatial transcriptomics methodologies. This comprehensive dataset serves as a valuable resource for future studies, enabling researchers to explore positional information, tissue remodeling, and the regulation of regeneration in biological systems.

## Materials and Methods

### Animal culture

Asexual *Schmidtea mediterranea* (strain CIW4) were maintained at 20 °C in a recirculating 1X Montjuic salts solution without antibiotics, following a previously described protocol (*68*). The animals were routinely fed beef liver. For experimental procedures, the animals were transferred to static culture and fasted for at least 7 days.

### Gene cloning and RNAi feeding

Genes of interest were cloned from a CIW4 cDNA library into the pPR-T4P vector as previously described (*69*). The resulting plasmids were used to produce dsRNA for RNA interference (RNAi). RNAi was performed following established protocols for gene knockdown (*70*). Briefly, bacterial pellets expressing the dsRNA were mixed with fresh beef liver paste in a 4:1 ratio. EGFP dsRNA was used as a control. Animals were fed every 3 days for a total of four or six RNAi feedings. After the final RNAi feeding, animals were amputated 3 days later to collect samples at various stages of regeneration.

### *In situ* hybridizations

Whole-mount in situ hybridizations were conducted following previously established protocols (*29, 71*). In short, the mucus from the animals was removed using 5% NAC in PBS, and then fixed for 1 hour in 4% formaldehyde (FA) in PBSTx (0.5%). The animals were bleached with formamide and incubated with proteinase K (2 μg/mL, AM2546, ThermoFisher) for 10 minutes. After a 2-hour pre-hybridization step, the hybridization was performed at 56°C for over 16 hours. Following extensive washes, the antibody signal was amplified using the Tyramide Signal Amplification system. Tissue clearing was achieved using ScaleA2 to reduce background noise (*29*).

### Sample fixation and section preparation for Stereo-seq

Sample fixation was carried out using a modified version of a previously described protocol (*72*). In short, animals were relaxed in 0.66 M MgCl₂ for 1 minute, followed by fixation in Methacarn solution (6 mL methanol, 3 mL chloroform, 1 mL glacial acetic acid) for 10 minutes. After fixation, the animals were rinsed in methanol three times, rehydrated in 50% methanol in PBS for 5 minutes, and then dehydrated in 20% sucrose in PBS for two cycles. The dehydrated tissues were embedded in pre-cooled OCT, frozen with dry ice, and stored at -80 °C until cryosectioning. Tissues were equilibrated in a -20 °C freezing microtome for 30 minutes prior to sectioning. RNA quality from cryosections was assessed using an Agilent 2100 Bioanalyzer. The cryosections of *Schmidtea mediterranea* were cut serially at 10 μm intervals using a Leica CM1950 cryostat. Each section was placed onto a Stereo-seq chip, incubated for 3 minutes at 37 °C on a Thermocycler Adaptor, and then fixed in methanol at -20 °C for 40 minutes.

### ssDNA staining and imaging of Stereo-seq slides

Prior to tissue permeabilization, sections on the Stereo-seq chip were stained with a nucleic acid dye (Thermo Fisher, Q10212) to visualize single-stranded DNA (ssDNA). The stained sections were then imaged using a Leica DM6M microscope. The images were stitched together and processed using the Leica Application Suite X software.

### Library construction and sequencing of Stereo-seq data

The library construction and sequencing protocols for Stereo-seq have been previously described (*27*). In summary, tissue sections were first washed with 100 μL of 0.1× saline-sodium citrate buffer (SSC, Thermo, AM9770) containing 0.05 U/μL RNase inhibitor (NEB, M0314L) to remove any remaining staining solution. Sections were then permeabilized using 0.1% pepsin (Sigma, P7000) in 0.01 M HCl buffer (pH 2.0) and incubated at 37 °C for 18 minutes. Released mRNAs were captured on the Stereo-seq chip and reverse transcribed overnight at 42 °C using SuperScript II reverse transcription mix (Invitrogen, 18064-014), containing 10 U/μL reverse transcriptase, 1 mM dNTPs, 1 M betaine solution, 7.5 mM MgCl₂, 5 mM DTT, 2 U/μL RNase inhibitor, 2.5 μM Stereo-seq template switch oligo, and 1× First-Strand buffer.

After in situ reverse transcription (RT), tissue sections were treated with a removal buffer (10 mM Tris-HCl, 25 mM EDTA, 100 mM NaCl, 0.5% SDS) at 37 °C for 30 minutes. The remaining RT products were then collected and amplified using KAPA HiFi Hotstart Ready Mix (Roche, KK2602) and 0.8 μM cDNA-PCR primers. PCR products were used to prepare sequencing libraries, with the following steps: quantification of concentration using the Qubit™ dsDNA Assay Kit (Thermo, Q32854), DNA fragmentation with in-house Tn5 transposase at 55 °C for 10 minutes, PCR amplification (KAPA HiFi Hotstart Ready Mix, Roche, KK2602) with 0.8 μM cDNA-PCR primers, and purification using Vazyme (N411-03). The purified PCR products were used to construct DNB libraries and sequenced on an MGI DNBSEQ-T1 sequencer (35 bp for Read1, 100 bp for Read2). The sequencing data were processed to generate a quantified spatial gene expression matrix at the subcellular level.

### Spatial transcriptomics data processing

Spatially resolved single-cell RNA-seq data obtained through Stereo-seq were pre-processed for further analysis. The first read (Read1) of the sequencing library contained coordinate identifiers (CIDs), molecular identifiers (MIDs), and poly-T sequences, while the second read (Read2) provided the captured cDNA sequences. The x-y coordinates of cDNA (715 nm resolution) were determined based on the CID sequences with a 1-bp mismatch tolerance. cDNA sequences were aligned to the *S. mediterranea* genome (dd_Smes_G4), and only mapped reads were used to identify exon transcripts. The MID sequences served to provide unique molecular identifiers (UMIs) for transcript quantification, with PCR duplicates removed using handleBam (https://github.com/BGIResearch/handleBam). Read pairs with a MID quality score below 10 were excluded. Finally, gene expression matrices incorporating spatial information were generated using clean exonic data.

### 3D reconstruction, clustering and cell type annotation of regenerative animals

Regenerating animals were reconstructed using methods outlined in an accompanying manuscript, where we developed a 3D spatial transcriptomics framework (Han et al., submitted). First, the MIRROR algorithm was applied to align the gene expression heatmap with the ssDNA staining image. Cell segmentation was then carried out using CellProfiler and Fiji. Gene expression data were mapped to each individual cell, creating a spatial transcriptome map at single-cell resolution. Next, after performing dimensionality reduction and clustering, cell clusters were annotated based on known lineage markers. The SEAM algorithm was employed to align the sections along the z-axis, thus determining the x-y-z coordinates of each cell. Morphological changes in the animals, caused by experimental procedures, were corrected based on established polarity gene patterns, and the 3D reconstructions were created using a combination of 3DSlicer and MeshLab. Finally, SPC cells from different stages of regeneration were integrated using the FindIntegration and IntegrateData functions in Seurat (v4.0.2). Dimensionality reduction and clustering were then conducted in Seurat following standard procedures.

### Blastema region detection in 3D spatial transcriptomics data

The blastema region was identified and segmented in the spatial coordinates of the reconstructed 3D model based on the pigmentation patterns observed in planarian body images. The boundaries separating the blastema from the rest of the body were defined using differences in pigmentation, which were quantified using the Threshold function in ImageJ. A quadratic polynomial regression was applied to refine these boundaries. The microscopy image and the corresponding 2D projection of the 3D model along the A/P-M/L (x, y) axis were aligned using TrakEM2. Each cell was then labeled with its corresponding region based on its position within the defined boundaries.

### Spatial pattern clustering of regenerating animals

To assess the expression dynamics within each cluster defined in homeostasis (Han et al., submitted), gene expression data from different stages of regeneration were mapped onto the appropriate clusters using linear regression. The regenerated animals were then divided into 100 bins along the A/P axis, in line with the homeostatic reference, and gene expression was categorized for each sample separately.

### Application of Turing pattern models to SBGs

We hypothesize that the interactions between SBGs and their upstream regulators follow Turing patterns within an autoregulatory activator-inhibitor framework. After removing the influence of spatial gradients, temporal gene expression data across eight regenerative time points were normalized, interpolated, and smoothed. Reaction, degradation, and diffusion parameters were fine-tuned using linear regression. These optimized parameters allowed us to predict gene expression levels at any post-amputation time point for both known and potential PCGs candidates. Pearson’s correlation coefficients were then calculated to assess the accuracy of these predictions.

### Principal Component Analysis (PCA) of SBGs and PCGs

To integrate temporal changes in gene expression with spatial variations, PCA was applied to the binned expression data of selected PCGs along the three axes in homeostatic animals (Han et al., submitted). For the A/P axis, the training set included 25 known PCGs. The first principal component (PC1), which accounted for 64.7% of the variance, corresponded to the head-tail gradient, while the second principal component (PC2), which explained 24.7% of the variance, captured fluctuations in the pharyngeal region (convex and concave). Based on their locations in the reduced-dimensional space, genes were manually grouped into five categories: head, Head-pharynx, Trunk-pharynx, pharynx-tail, and tail domains.

Because there were few previously identified PCGs with clear spatial patterns along the M/L and D/V axes in our Stereo-seq data, we expanded the training sets to include 42 and 67 newly inferred PCGs, respectively. Potential M/L PCGs candidates were selected based on Spearman’s rank correlation coefficients greater than 0.7 or less than -0.7 for binned SCT-transformed gene expression data, showing patterns similar to known PCGs. For potential D/V PCGs, a fold change greater than 1.5 between dorsal and ventral regions in binned SCT-transformed values was used as a selection criterion.

PCA was performed using the Scikit-learn package with default parameters. The eigenvectors derived from this analysis were used to map both known and potential PCGs candidates back into the reduced PCA space, trained on the homeostatic data. The regenerative trajectories of PCGs were visualized, resembling the behavior of an underdamped mass-spring system: the initial amputation stretched the “spring,” disrupting the PCG expression profile, while the gene regulatory network provided feedback to restore the profile to its homeostatic state.

### UV unwrapping for head blastema epidermis region

To convert the 3D body shell of the planarian into a 2D plane, we employed the UV unwrapping technique, a common procedure in the field of computer graphics, using Blender (https://www.blender.org/), an open-source 3D creation suite. UV unwrapping is a process in which the surface of a 3D model is mathematically “unwrapped” and mapped onto a 2D plane, enabling precise application of textures and structures onto a flat surface, which is essential for accurate visualization and analysis. The process of UV unwrapping for the head blastema epidermis involved segmenting the planarian body, marking seams for accurate texture alignment, unwrapping the mesh, and mapping the epidermal cells onto a 2D plane. This approach enabled a high-precision representation of the head blastema’s epidermal region, which will be useful for further texture analysis and studies related to planarian regeneration.

Briefly, the planarian body mesh was first segmented into two sections along a defined plane located near the boundary of the head blastema. This segmentation was performed using the Bisect Tool within Blender, which allowed for a clean division of the mesh without distorting the geometry. The separation enabled us to isolate the head blastema area, which we intended to unwrap for detailed analysis.

Next, to prepare for the unwrapping process, seams were strategically marked to guide the unfolding of the 3D mesh. The edges connecting the blastema cutting plane to the anterior pole of the planarian were designated as a seam, specifically placed along the D/V boundary. This step ensures that the unwrapping process adheres to natural anatomical divisions, preventing distortion of the texture in the subsequent 2D plane. The seam marking was carried out using Blender’s UV editor tool, where the mesh’s geometry was manipulated to set boundaries for the unwrapping operation.

Once the seams were defined, the head blastema mesh was subjected to the UV unwrapping operation. The mesh was unfolded into a 2D plane, with particular attention paid to the correct alignment of the marked seam. The seam was clipped and adjusted to create a smooth, curved incision, which represents the epidermal cells of the head blastema accurately in flat space. This incision helped ensure that the texture mapping would preserve the anatomical integrity of the original 3D model.

Finally, we focused on the outer epidermal cells of the head blastema, which were part of the original 3D point cloud data. These epidermal cells were mapped onto the unwrapped 2D plane using a process that minimized the distance between the original 3D coordinates of each epidermal cell and the vertices of the subdivided mesh surface. This step ensured a precise mapping of cellular structures to the 2D plane, allowing for high fidelity in representing the epidermal region’s texture and topology. The minimization of distances between the original 3D points and the 2D unwrapped model helped preserve spatial relationships and accurately represented the cellular organization in a flattened format.

### Monocle3 analysis

To investigate the state transitions of the ARZ (Clu.31 cluster) during regeneration, we analyzed the trajectory dynamics using raw transcriptome counts from 36 hours post-amputation (hpa) to 14 days post-amputation (dpa). The raw counts were first normalized using SCTransform in Seurat (v4.0.2). Dimensionality reduction and clustering were then performed in Monocle3 (v1.3.1) (*41*). he trajectory graph was constructed by fitting the principal graph with the learn_graph function, and pseudotime was calculated with the neoblast cell type set as the root. Marker genes for the ARZ (Clu.31 cluster) were identified using Seurat, and their expression was projected onto the trajectory branches. Cells were clustered in a 15-NN graph, which allowed us to divide the trajectory into distinct branches enriched for epidermal, muscle, and neuronal signatures.

### RNA velocity analysis

RNA velocity analysis, based on Waddington’s epigenetic landscape and differential geometry, was used to make continuous, time-resolved predictions of cell state transitions. Cellular genes were aligned to the reference genome to identify exon and intron sequences. The relative abundance of spliced (mature) and unspliced (nascent) mRNAs was calculated to estimate splicing and degradation rates using Velocyto (*42*). E Each DNB was assigned to its corresponding cell based on its x and y coordinates. Spliced and unspliced count matrices for different cell types were processed using the recipe_monocle function in Dynamo (*73*) to identify highly expressed genes. Following dimensionality reduction, the continuous velocity vector field was reconstructed in UMAP space to predict future cell fates.

### Monocle2 analysis

Monocle2 (v2.18.0) was used to analyze the ventral epidermal trajectory across different regeneration time points, following the tutorial (http://cole-trapnell-lab.github.io/monocle-release) (*74*).We focused on extracting SPC clusters during the putative ventral epidermal transition from the blastema region. Differentially expressed genes for each cell type were identified and used to order the cells. Dimensionality reduction was carried out using the DDRTree method, and the plot_cell_trajectory function was used for visualization. Marker genes identified by Seurat’s FindAllMarkers function were projected along the estimated pseudotime to assess their potential role in the transition.

### Visual representation of the website: Planarian Regenerative Interactive Spatiotemporal Transcriptomic Atlas in Four Dimensions (PRISTA4D)

To enhance the accessibility and utility of our 4D planarian cell atlas for researchers in the field of regeneration, we developed an open-source, interactive database (available at https://www.bgiocean.com/planarian). This platform enables users to explore the spatial distribution and dynamic changes of various genes and cells at different stages of regeneration.

The PRISTA4D database provides several functions, including browsing capabilities and access to experimental procedures, data analysis pipelines, and the ability to download the original dataset. It serves as an invaluable resource for studying cell differentiation and spatiotemporal cell interactions within the regeneration research community.

The website includes a homepage and five key functional modules:

3D Model: Visualizes different cell types in three dimensions, allowing users to view cellular organization across regeneration stages.

Spatial Clustering Module: Illustrates the distribution of genes and cell types, offering insights into spatial gene expression patterns.

Stereo-seq Module: Provides detailed experimental protocols used to generate the transcriptomic data, ensuring transparency and reproducibility.

Sampling Design Module: Offers information on sample design and the data analysis pipelines used, enabling users to understand how the data was processed and analyzed.

Download Module: Grants access to the complete original dataset, allowing users to download the raw data for further analysis and research.

### Cell sorting and library construction for single-cell RNA sequencing

To prepare the cell suspension for scRNA-seq, CMFB buffer (CMF + 1% FBS) was placed on a cold plate (4 °C), and the animals were incubated in this solution before their tissues were manually chopped to release cells as described previously (*8*). After dissociation, the cells were pelleted by centrifugation at 290 g for 5 minutes at 4 °C. The suspension was filtered through a 40 μm filter and stained with DAPI (1:1000; Beyotime) and DRAQ5 (1:1000; BioLegend). The cells were washed and resuspended in CMFB buffer. Flow cytometry and sorting were performed using a Sony MA900 cell sorter, with the temperature kept at 4 °C to maintain cell integrity. Around 20,000 viable cells (DAPI-; DRAQ5+) were loaded onto the SeekGene platform using the Single Cell 3’ Transcriptome kit to generate scRNA-seq libraries. Library preparation followed the manufacturer’s guidelines to ensure optimal coverage, and sequencing was conducted on an Illumina NovaSeq platform with paired-end 150 base pair (150 PE) reads for detailed transcriptomic profiling.

### Analysis of single-cell RNA sequence data after RNAi knockdown

Raw scRNA-seq data were processed and aligned with the *S. mediterranea* reference transcriptome (smed_20140614). Cells that had fewer than 200 detected features or genes expressed in fewer than three cells were excluded. Cells exhibiting unusually high mitochondrial or ribosomal RNA content (where mitochondrial percentage was more than twice the median) were also filtered out based on the annotated reads (data S6). After quality control, the remaining Unique Molecular Identifiers (UMIs) were quantified and analyzed using the Seurat package in R, which enabled normalization, scaling, and dimensionality reduction of the data. Principal component analysis (PCA) was performed, and the top 30 principal components were used for 2D UMAP generation and clustering within Seurat. Cell lineages were annotated based on the expression of known marker genes, while SPC clusters were identified using markers derived from 3D spatial transcriptomics. Batch effects due to technical variations across samples were corrected using Seurat’s integration functions, ensuring that the observed differences were biologically relevant. For analyzing cellular differentiation trajectories and lineage relationships, Monocle 2 was applied. Additionally, MiloR was used to examine and compare the abundance of cells in specific neighborhoods or microenvironments between the control and RNAi-treated groups (*75*). Briefly, PCA was used for dimensionality reduction, and a KNN graph was constructed with the buildGraph function (k=30, d=30) based on the top 30 PCA dimensions. Neighborhoods were defined using the makeNhoods function (prop=0.1, k=30, d=30), and differential abundance testing was conducted with default parameters using the distinct function. Differentially abundant cell populations between control and knockdown groups were identified, and differentially expressed genes (DEGs) were determined using DESingle (v1.9.2) with standard settings (*76*). Genes with an adjusted p-value of less than 0.05 were considered significantly different. This comparative analysis revealed how RNAi interventions affect the cellular composition within the tissue.

## Acknowledgments

We thank L. Bolund, D. Little and all Zeng lab members for the critical reading of the manuscript.

## Funding

This research was supported by the National Key R&D Program of China (2022YFC3400400), the National Key R&D Program of China (2020YFA0112502 and 2021YFA1100202 to A.Z.), the National Natural Science Foundation of China (32070828 to A.Z.), Shenzhen Science and Technology Program (JCYJ20250604191305008 to M.X. and RCJC20221008092804002 to Y.G.), the Strategic Priority Research Program of the Chinese Academy of Sciences (XDA16021300), the CAS Pioneer Hundred Talents Program (A.Z.), Shanghai Pujiang Program (20PJ1414600 to A.Z.), the Shanghai Science and Technology Committee (STCSM) (22ZR1468400 to A.Z.), Guangdong Genomics Data Center (2021B1212100001) and the Feng Foundation of Biomedical Research (A.Z.).

## Author contributions

A.Z., X.X., G.F., H.L., K.H. and M.X. conceived and directed the study. A.Z., X.X., M.X. and H.L. supervised the work. Y.C. and YR.L. performed animal experiments and RNAi. XW.L., W.G., JQ.W., W.W. and H.P. performed the Stereo-seq experiments. K.H., M.X., L.G., Y.L., Y.W. and Z.H. analyzed the data. L.G., Y.L. and T.Y. performed database construction. Q.L., L.Z. and X.M. assisted in the data analysis. R.Z., L.L., X.W., H.Z., X.S., S.L., W.Z., S.T.C, J.F., X.L., Y.G., J.W. G. P., and H.Y. performed investigations. A.Z., G.F., H.L., M.X. and K.H. wrote the manuscript with input from all the authors.

## Competing interests

Authors declare that they have no competing interests.

## Data and materials availability

All data generated in this study were deposited at CNGB Nucleotide Sequence Archive (accession code: STT0000028). The accession number for Rod1 (SMED30003831) is OR211556. Processed data can be interactively explored from our PRISTA4D database (https://db.cngb.org/stomics/prista4d) or 3D modeling website: https://www.bgiocean.com/planarian). All original codes supporting the current study are hosted on GitHub (https://github.com/BGI-Qingdao/4D-BioReconX). The web-based 3D interactive PRISTA4D database has also been made available (https://www.bgiocean.com/planarian). Any additional information required to reanalyze the data reported in this paper is available from the lead contact upon request.

## Notes

### Competing Interest Statement

The authors have declared no competing interest.

