## Supplementary Figures for "4D Single-Cell Spatial Transcriptomics Reveals Dynamic Morphogenetic Gradients and Regenerative Domains in Planarians"

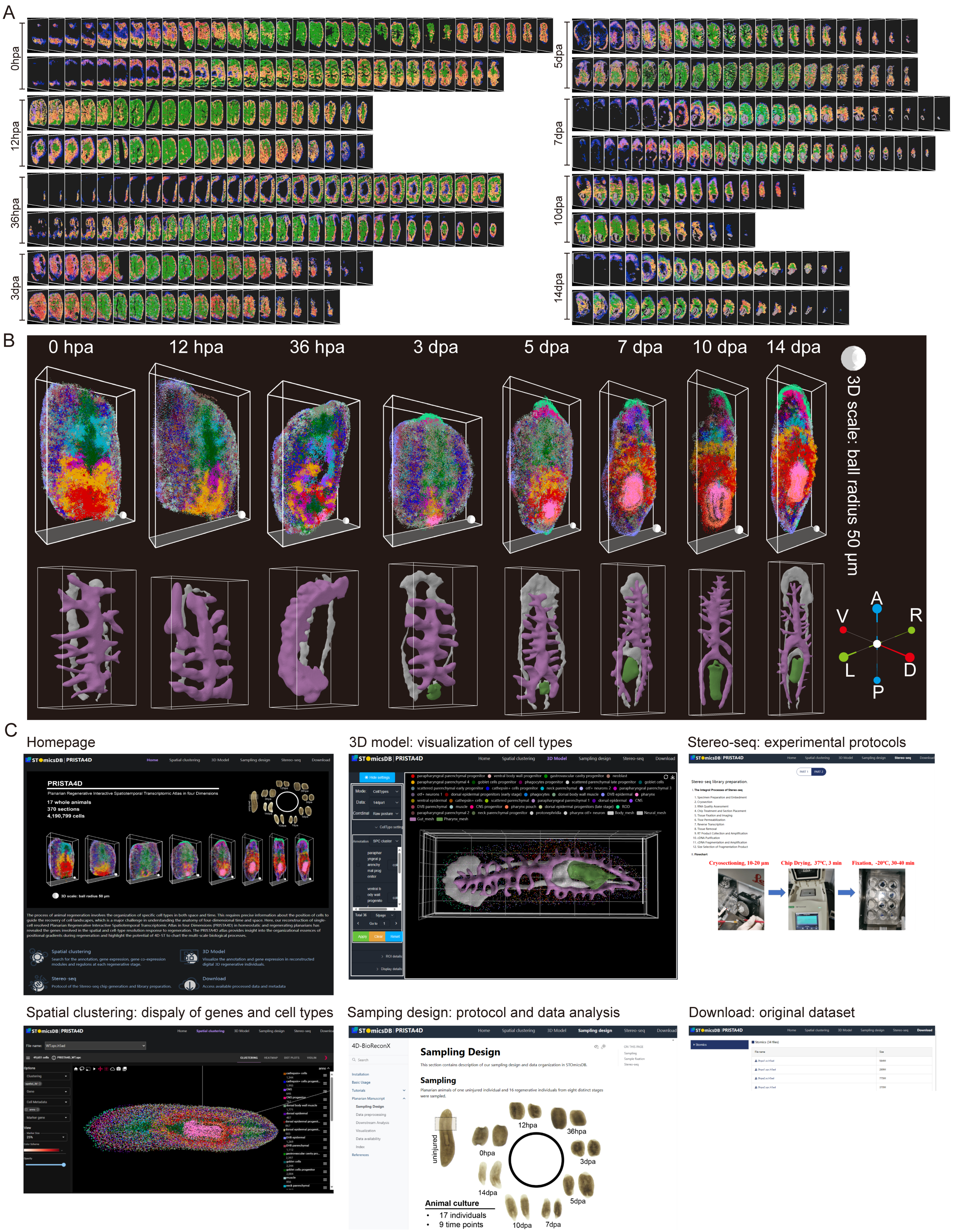
A
0hpa
12hpa
36hpa
3dpa
5dpa
7dpa
10dpa
14dpa
B
0 hpa
12 hpa
36 hpa
3 dpa
5 dpa
7 dpa
10 dpa
14 dpa
3D scale: ball radius 50 μm
A
V
R
L
D
P
C
Homepage
3D model: visualization of cell types
Stereo-seq: experimental protocols
Spatial clustering: dispaly of genes and cell types
Samping design: protocol and data analysis
Download: original dataset
PRISTA4D
17 whole animals
370 sections
4,190,799 cells
Cryosectioning, 10-20 μm
Chip Drying, 37°C, 3 min
Fixation, -20°C, 30-40 min
Sampling Design
Sampling
uninjured
0hpa
12hpa
36hpa
3dpa
5dpa
7dpa
10dpa
14dpa
Animal culture
17 individuals
9 time points

**Fig. S1. High-resolution 4D spatial and molecular characterization of planarian regeneration.**
**(A)** Consecutive Stereo-seq sections used for 3D reconstruction across eight regenerative time points. Two animals were sampled per time point, with each row representing one animal. Sections display the spatial distribution of cell types from ventral (V) to dorsal (D), moving left to right. Cells are color-coded according to lineage annotations as shown in Fig. 1B. hpa, hours post-amputation; dpa, days post-amputation.
**(B)** 3D spatial visualization of 36 SPC clusters (top row) and tissue meshes (bottom row) across the same eight regenerative time points. Cells are color-coded based on their SPC cluster annotations, as shown in Fig. 1B. The anterior of the worm is oriented upwards, with a dorsal view shown.
**(C)** Overview of the interactive PRISTA4D (Planarian Regenerative Interactive Spatiotemporal Transcriptomic Atlas in Four Dimensions) website (https://db.cngb.org/stomics/prista4d/). The homepage and five key functional modules are highlighted. The 3D model visualizes cell types, while the spatial clustering module illustrates the distribution of genes and cell types. The Stereo-seq module details experimental protocols, and the sampling design module provides information on sample design and data analysis pipelines. The download module grants access to the full original dataset.

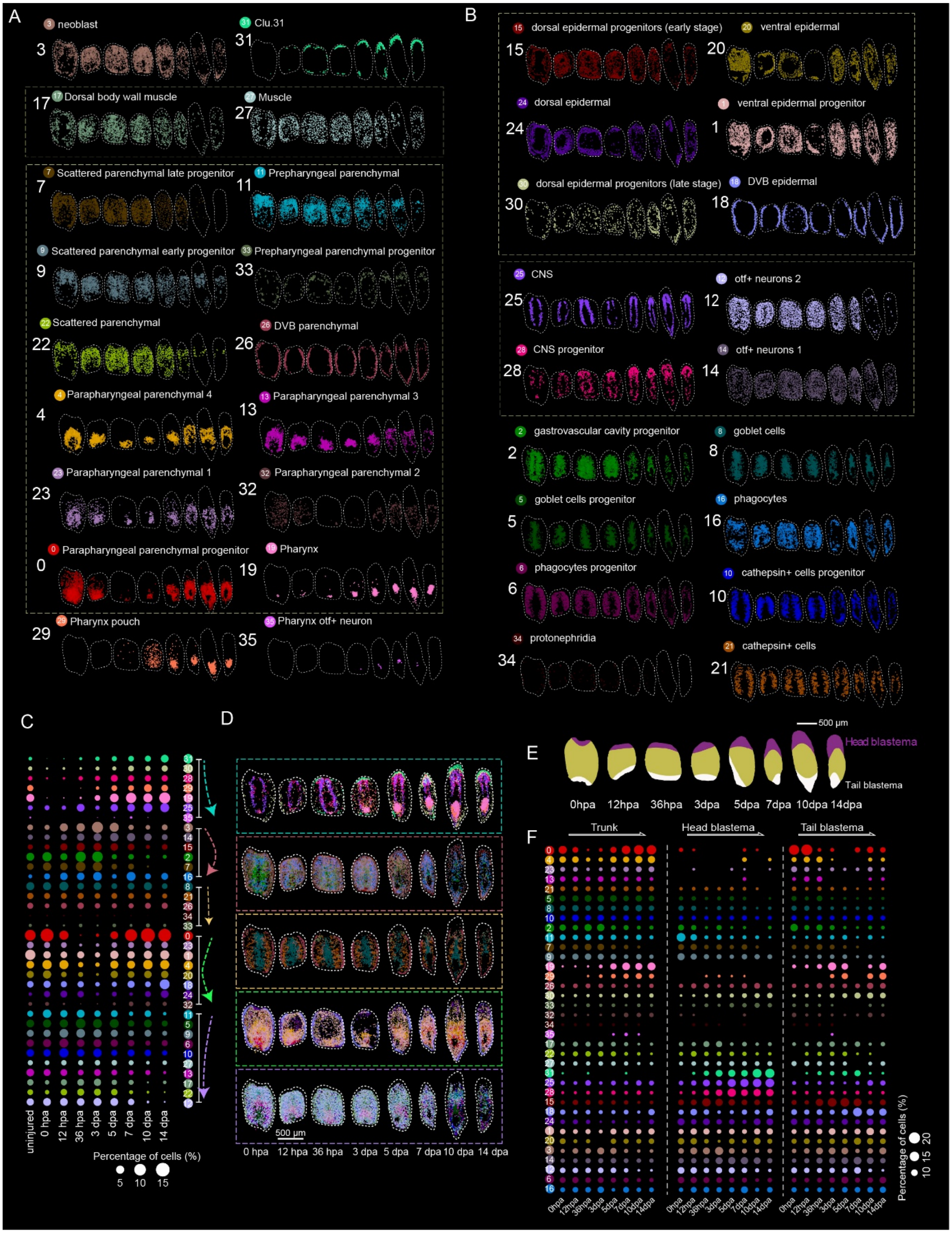
A
3 neoblast
31 Clu.31
17 Dorsal body wall muscle
27 Muscle
7 Scattered parenchymal late progenitor
11 Prepharyngeal parenchymal
9 Scattered parenchymal early progenitor
33 Prepharyngeal parenchymal progenitor
22 Scattered parenchymal
26 DVB parenchymal
4 Parapharyngeal parenchymal 4
13 Parapharyngeal parenchymal 3
23 Parapharyngeal parenchymal 1
32 Parapharyngeal parenchymal 2
0 Parapharyngeal parenchymal progenitor
19 Pharynx
29 Pharynx pouch
35 Pharynx otf+ neuron
B
15 dorsal epidermal progenitors (early stage)
20 ventral epidermal
24 dorsal epidermal
1 ventral epidermal progenitor
30 dorsal epidermal progenitors (late stage)
18 DVB epidermal
25 CNS
12 otf+ neurons 2
28 CNS progenitor
14 otf+ neurons 1
2 gastrovascular cavity progenitor
8 goblet cells
5 goblet cells progenitor
16 phagocytes
6 phagocytes progenitor
10 cathepsin+ cells progenitor
34 protonephridia
21 cathepsin+ cells
C
uninjured
0 hpa
12 hpa
36 hpa
3 dpa
5 dpa
7 dpa
10 dpa
14 dpa
Percentage of cells (%)
5
10
15
D
500 μm
0 hpa
12 hpa
36 hpa
3 dpa
5 dpa
7 dpa
10 dpa
14 dpa
E
500 μm
Head blastema
Tail blastema
0hpa
12hpa
36hpa
3dpa
5dpa
7dpa
10dpa
14dpa
F
Trunk
Head blastema
Tail blastema
0hpa
12hpa
36hpa
3dpa
5dpa
7dpa
10dpa
14dpa
Percentage of cells (%)
10
15
20

**Fig. S2. Characterization and visualization of cellular populations across regeneration timepoints.**

**(A)** Spatial visualization of 18 SPC clusters at eight regeneration time points, including neoblasts, the Clu.31 domain, muscle lineages (2 subtypes), parenchymal cells (11 subtypes), and pharyngeal cells (3 subtypes).

**(B)** Spatial visualization of 18 SPC clusters at eight regeneration time points, including epidermal cells (6 subtypes), neuron lineages (4 subtypes), intestinal cells (5 subtypes), cathepsin-expressing cells (2 subtypes), and protonephridia lineages.

**(C)** Bubble plots illustrating changes in cell ratios of SPC clusters at uninjured and eight regenerative time points. SPC clusters are grouped by population dynamics, with trends indicated by arrows. Cluster numbers are shown on the right, with numbered circles.

**(D)** 3D projections of spatially grouped SPC clusters at eight regenerative time points. Cluster groups correspond to those indicated in the bubble plots. Scale bar, 500 µm.

**(E)** Animatic visualization of the head blastema (purple), trunk (brown), and tail blastema (white) regions in each regenerating animal. The blastema regions are identified based on pigmentation. Scale bar, 500 µm.

**(F)** Bubble plot showing changes in the ratio of SPC clusters across eight regenerative time points in the trunk, head, or tail blastema regions.

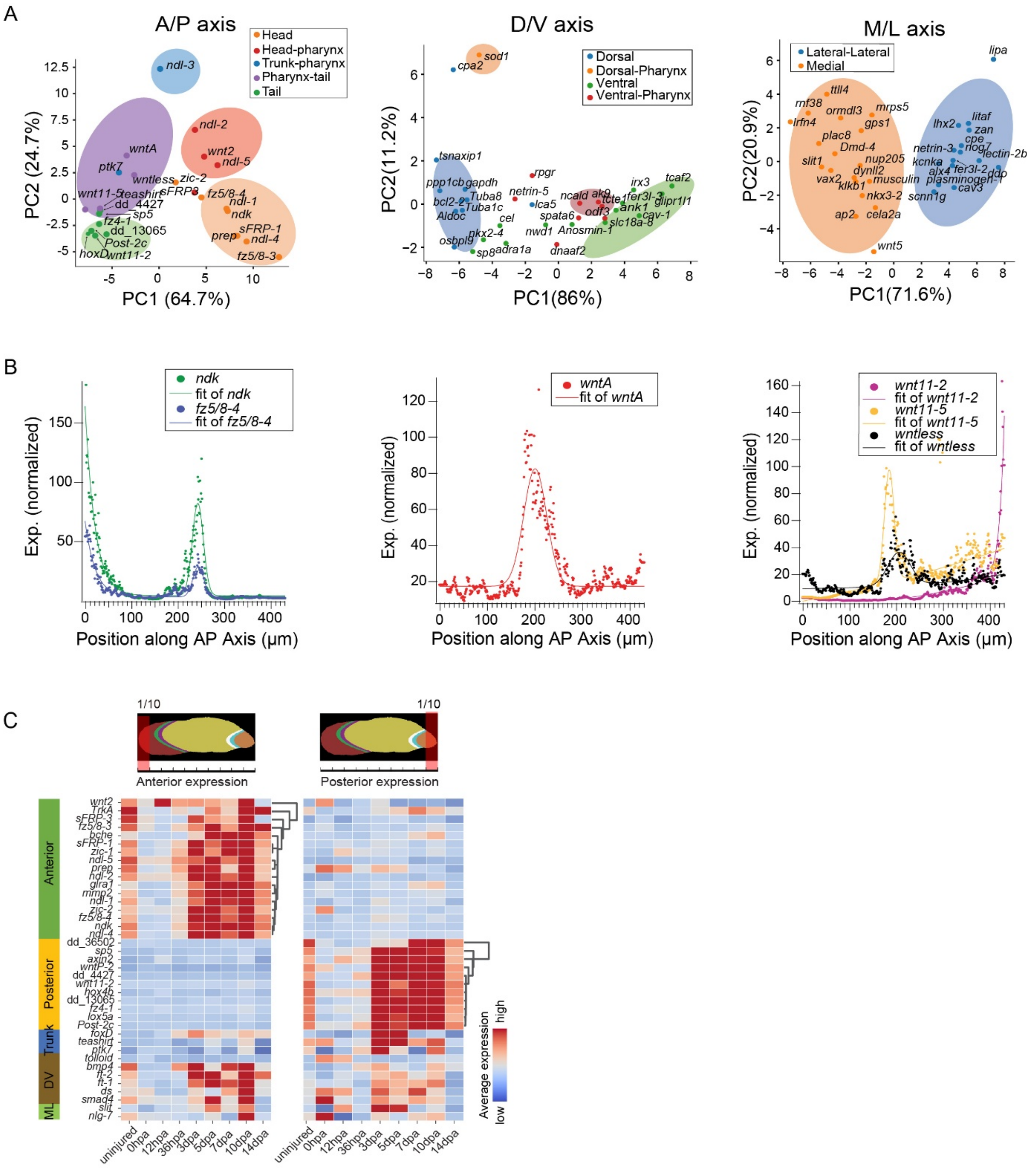
A
A/P axis
Head
Head-pharynx
Trunk-pharynx
Pharynx-tail
Tail
PC1 (64.7%)
PC2 (24.7%)
D/V axis
Dorsal
Dorsal-Pharynx
Ventral
Ventral-Pharynx
PC1(86%)
PC2(11.2%)
M/L axis
Lateral-Lateral
Medial
PC1(71.6%)
PC2(20.9%)
B
ndk
fit of ndk
fz5/8-4
fit of fz5/8-4
wntA
fit of wntA
wnt11-2
fit of wnt11-2
wnt11-5
fit of wnt11-5
wntless
fit of wntless
Exp. (normalized)
Position along AP Axis (μm)
C
1/10
Anterior expression
Posterior expression
Anterior
Posterior
Trunk
DV
ML
Average expression
high
low
uninjured
0hpa
12hpa
36hpa
3dpa
5dpa
7dpa
10dpa
14dpa

**Fig. S3. Self-organized gradient formation during polarity regeneration.**

**(A)** PCA representations of gene expression patterns for A/P (left), M/L (middle), and D/V (right) PCGs or PCG candidates in intact animals. Variance explained by principal components (PC1: 64.7%, PC2: 24.7% for A/P axis; PC1: 86.0%, PC2: 11.2% for D/V axis; PC1: 71.6%, PC2: 20.9% for M/L axis).

**(B)** Fitting plots of PCG spatial patterns to an exponential function (with optional Gaussian fit for the pharynx) for anterior-enriched (left), pharynx-enriched (middle), and posterior-enriched (right) PCGs in intact animals.

**(C)** Heatmaps displaying the dynamic expression levels of indicated genes in the anterior or posterior regions of the body (1/10 body length) during different timepoints of regeneration. Genes showing enrichment in specific regions, with similar temporal patterns, are arranged by hierarchical clustering as shown on the left.

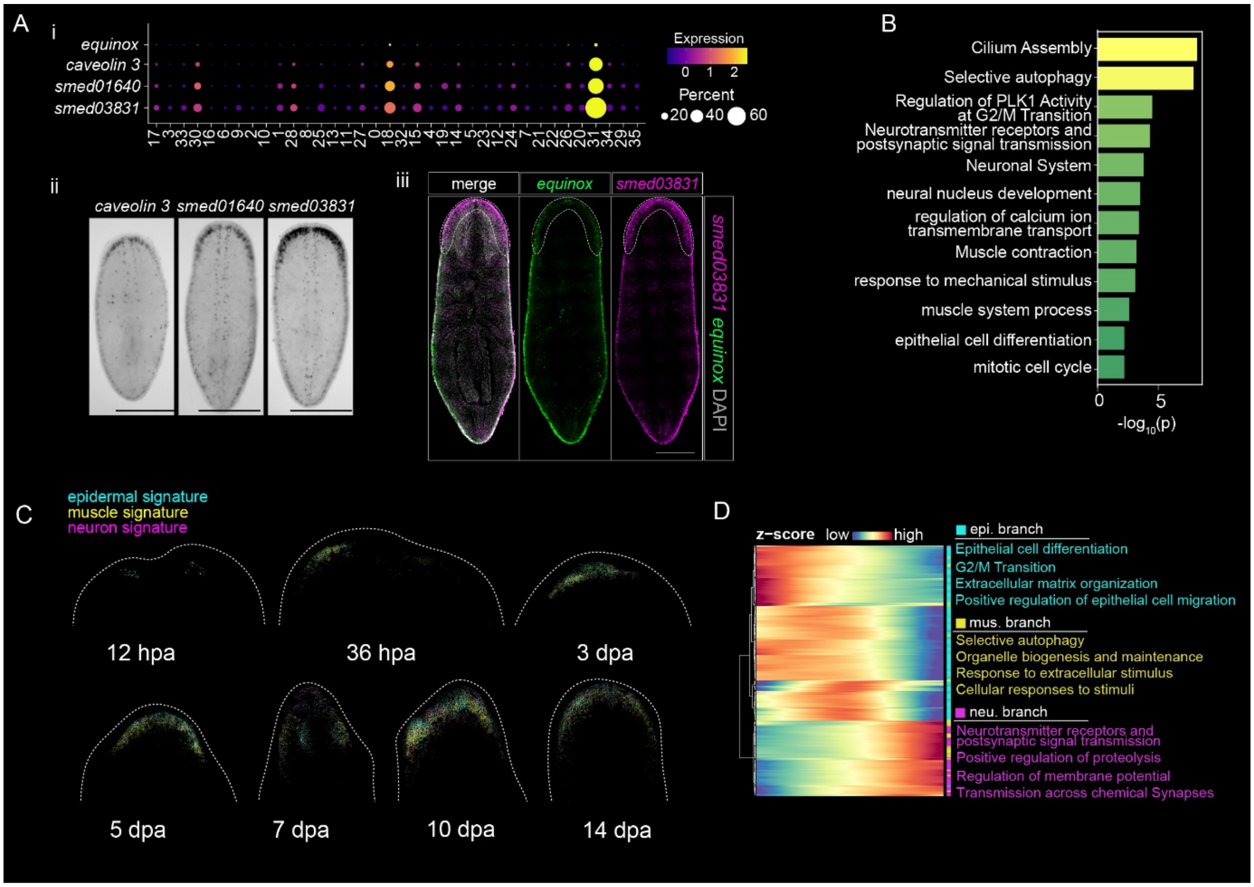


**Fig. S4. Composition and molecular signatures of the ARZ (Clu.31) domain.**

**(A)** Characterization of the Clu.31 domain. (i) Bubble plot showing the expression of the known wound epidermis marker (equinox) and the top three Clu.31-enriched marker genes across all SPC clusters. (ii) Whole-mount in situ hybridization images of three Clu.31 markers in homeostatic animals. (iii) Double fluorescence in situ hybridization of a representative Clu.31 marker (*smed03831*) and *equinox*. Scale bars, 500 µm. n ≥ 3.

**(B)** Functional enrichment analysis of Clu.31-enriched genes in homeostatic animals.

**(C)** Spatial visualization of three lineages within the Clu.31 domain in the head blastema at different time points during regeneration.

**(D)** Heatmap displaying enriched gene expression along the Clu.31 trajectory, as shown in Fig. 3H. Functional enrichment terms for genes associated with each pseudotime branch are indicated on the right.

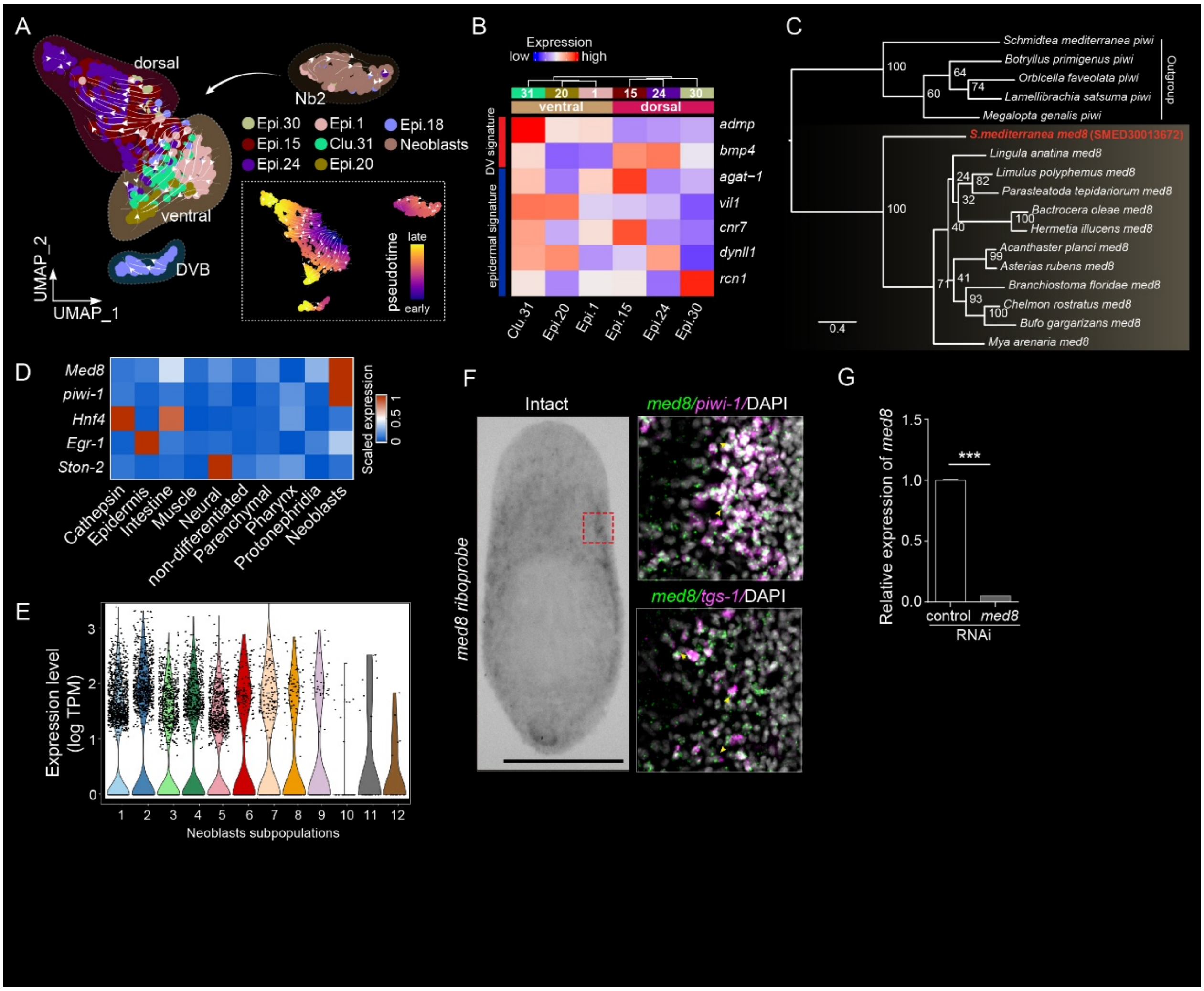


**Fig. S5. Characterization of the ARZ (Clu.31) domain and expression analysis of *med8*.**
**(A)** RNA velocity streamline plot illustrating the predicted cell transition trajectory at 36 hpa, when Clu.31 first emerges. Arrows indicate that Clu.31 primarily originates from Epi.1 (Cluster 1: ventral epidermal progenitor). Inset (bottom right): inferred pseudotime progression, represented by a color scale from earliest (blue) to latest (yellow).

**(B)** Heatmap showing the relative expression of epidermal signature genes and dorsal (*bmp4*) and ventral (*admp*) positional genes in Clu.31 and selected epidermal SPCs (Epi. 20, 1, 15, 24, and 30).

**(C)** Phylogenetic tree of Mediator 8 family proteins, with PIWI-1 serving as an outgroup.

**(D)** Heatmap displaying med8 expression across different cell populations. Data from Blair W. Benham-Pyle et al. (*17*).

**(E)** Violin plot showing med8 expression in different neoblast subpopulations. Data from An Zeng et al. (*8*).

**(F)** Whole-mount in situ hybridization (WISH) showing med8 expression (left) and co-expression with piwi-1 (top right) or tgs-1 (bottom right). n ≥ 6 animals showed a consistent pattern. Scale bar, 500 µm.

**(G)** med8 RNAi efficiency assessed by qRT-PCR.

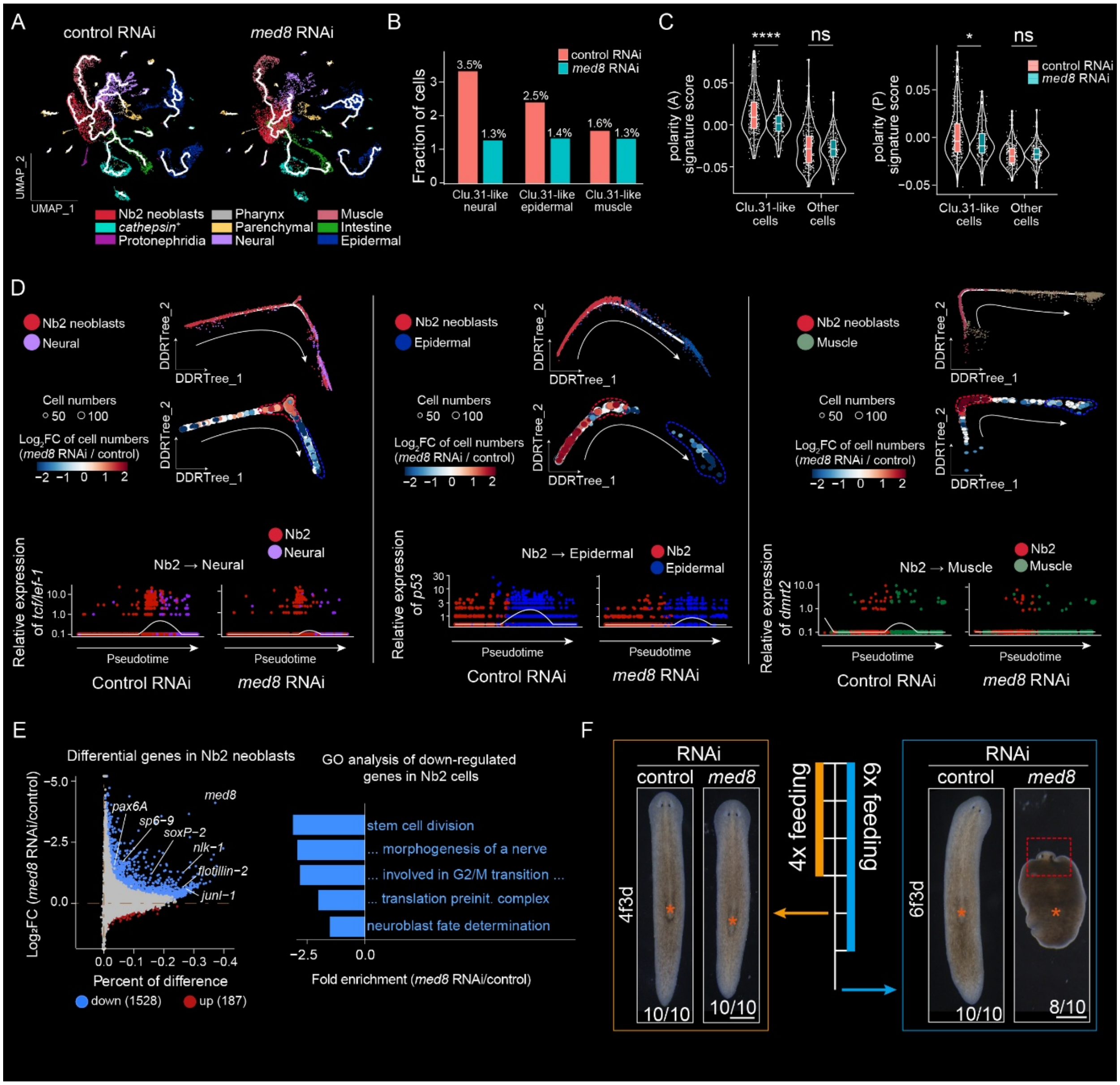


**Fig. S6. *med8*-regulated Clu.31 controls cell production during blastema formation.**

**(A)** Pseudotime trajectory analysis of scRNA-seq data showing potential impairment in the differentiation of Nb2 neoblasts into other cell lineages in med8 RNAi animals. Altered cell lineages are indicated by dashed lines.

**(B)** Bar plot comparing the fraction of lineage components within Clu.31 between control and med8 RNAi animals, based on scRNA-seq data.

**(C)** Violin plots displaying the signature scores of anterior (left) and posterior (right) polarity genes. Statistical significance was assessed using the Wilcoxon test (p). ns, $p > 0.05$; *, $p < 0.05$; ****, $p < 0.0001$.

**(D)** Pseudotime trajectory analysis using Monocle2 (top row) and differential cell abundance analysis with miloR (middle row) for predicted transitions from Nb2 neoblasts to neural (left), epidermal (middle), or muscle (right) lineages. Scatter plots (bottom row) show the relative expression of *tcf/lef-1*, *p53*, and *dmrt2* along the inferred Nb2 neoblast transition trajectories toward the indicated lineages.

**(E)** Left: Volcano plot displaying differentially expressed genes in Nb2 neoblasts following med8 RNAi. Right: Representative Gene Ontology enrichment terms for downregulated genes in Nb2 neoblasts after med8 RNAi.

**(F)** Representative phenotypes of *med8* RNAi animals after 4 or 6 feedings, observed 3 days after the last feeding. Scale bars, 500 μm.

**Supplemental Materials**

data S1. Summary statistics of Stereo-seq data.

data S2. Differentially expressed genes across regeneration time points.

data S3. Pattern clusters of positional control genes (PCGs) along the anterior-posterior (A/P) axis.

data S4. Cell type specificity of known and potential PCG candidates.

data S5. Signature gene list used for module score calculation.

data S6. List of ribosomal RNA (rRNA) and mitochondrial RNA (mtRNA) in *Schmidtea mediterranea*.
